# Analyzing Human Cytomegalovirus Genotype Diversity by Modelling Viral Population Dynamics

**DOI:** 10.64898/2025.12.29.696899

**Authors:** Raphael Eichhorn, Irene Görzer, Büsra Külekci, Madlen Mollik, Cornelia Pokalyuk

## Abstract

Some viruses including the widespread human cytomegalovirus (HCMV) show a high level of genetic diversity within hosts and across the whole virus population even in relatively conserved genomic regions. To investigate how this diversity is formed, we earlier introduced and analyzed a population model for neutrally evolving viruses that persist in their hosts and are capable of reinfection, mutation and recombination. Based on these results, we fit our model to observed genotype frequencies from Austrian HCMV patients. Despite simply assuming neutral evolution, the model captures the data closely. The inferred parameters are discussed and may give insights about the roles of viral replication, reinfection, mutation and recombination for the evolution of HCMV.

---

The genetic diversity observed in viruses is the result of an intricate interplay of different evolutionary forces. In addition to mechanisms such as replication (or reproduction), mutation and recombination that govern the populations of many organisms, virus populations are subject to the structure, behavior and interactions of their host population. A priori, the effects of this setting on the dynamics of genetic diversity in viruses are not clear.

Understanding the origin of the genetic variability (within hosts) of a given virus is of great interest, since a highly variable within-host viral population may be harder for the host’s immune system to control. Different interventions may be relevant to suppress the accumulation of diversity depending on its origin. If, for example, de novo mutations generate genetic diversity, one would try to restrict population growth within a host. If, however, genetic diversity is constantly introduced in a host due to reinfection events, one might attempt to reduce contact with other infected hosts.

Viruses exhibit different degrees of variability at the population level. Among the human herpesviruses, which are restricted to their human host, *varicella zoster virus* (HHV-3) shows very little variation, with an overall mean distance of 0.0014 substitutions per site. This is in sharp contrast to the *human cytomegalovirus* (HCMV; HHV-5), which has an overall mean distance of at least 0.0266 [1] and possesses different genotypes at numerous loci [2, 3]. Furthermore, HCMV displays pronounced within-host genetic diversity, with hosts frequently being infected with multiple genotypes. HCMV is of sustained medical interest due to its high prevalence, with an estimated worldwide seroprevalence of 86% [4], and due to the significant morbidity it causes: It is *the* leading viral cause of birth defects, and while infections in healthy adults usually cause no or only mild symptoms, it can lead to severe disease and death in immunocompromised hosts like transplant recipients or HIV patients [5].

The causes of the emergence and maintenance of genetic diversity in HCMV remain largely unclear. Specifically the viral replication dynamics of different genotypes within single hosts are not fully understood. It is not clear if observed genotype frequencies within hosts and at the population level can potentially be attributed to different levels of fitness of the genotypes (depending on the host), to balancing selection or to random genetic drift. The first scenario seems rather unlikely, since in a relatively large proportion of hosts several genotypes can be detected at non-trivial frequencies [6, 7]. The scenario of balancing selection assumes a tendency towards an equilibrium frequency of genotypes within hosts. This could be reasonable if the presence of several genotypes within a host is beneficial for the virus because e.g. different immune responses can be circumvented more easily. This setting has been analyzed previously from a mathematical perspective [8]. Balancing selection has also been conjectured to contribute to HCMV diversity at some loci based on the analysis of 259 whole genome sequences [9]. The third scenario assumes equal fitness of all genotypes and has been studied mathematically before [10]. Here, the viral population is distributed (equally) over all infected hosts and as evolutionary factors mutation, viral replication, host reproduction, reinfection as well as recombination are considered.

Current quantitative research on virus populations, including HCMV, often focuses on a single time scale and on one or a few evolutionary effects at a time. Additionally, the hierarchical dependence on the host population, which virus populations are naturally subject to, is often neglected. With this work we aim to demonstrate how the stochastic model from [10] that incorporates effects across different time scales and a hierarchical structure between the host and the virus population, may be used to analyze genetic diversity in HCMV. We fit the mathematical model to genotype data. The thereby inferred parameters are discussed with respect to parameter estimates available in the literature, and give some insight about the roles of viral replication, reinfection, mutation and recombination for the evolution of HCMV.

While we concentrate on the application of the model to HCMV, the presented method is not limited to HCMV but may serve as a framework to be applied to other host-pathogen systems, in which the pathogens are capable of persistence and reinfection.

## 1 Model

We consider a population of *M* ∈ ℕ hosts that are all assumed to be infected with a certain virus and assume that the number of virus particles per host is constantly equal to *N* ∈ ℕ to model the persistent viral population. We are interested in (the dynamics of) the allele frequency spectrum at *L* ∈ ℕ loci. At locus *ℓ* the number of possible genotypes (or alleles) is *K*_*ℓ*_ ∈ ℕ for *ℓ* ∈ [*L*] where [*L*] := {1, …, *L*}. For simplicity of notation we assume that *K*_*ℓ*_ = *K* is the same for all loci but our results trivially can be generalized to the case of varying *K*_*ℓ*_. The type of a virus particle at the *L* studied loci is represented by a vector in [*K*]^*L*^ which we call the *sequence type* of that particle. We model the dynamics of the host-virus system by a Markov jump process **Z**^*M,N,L,K*^ taking values in

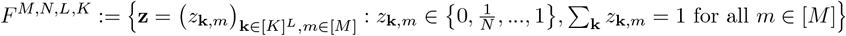

that tracks the relative frequencies of the *K*^*L*^ different sequence types in the *M* different hosts. We keep *L* and *K* fixed throughout the article and consider the joint limit of large host and virus population, hence we simply write **Z**^*N*^ and *F*^*N*^ in the following.

We assume that viral particles replicate, recombine, and mutate, as well as that hosts die and are replaced instantly by primary infected, that is so far uninfected, hosts. At these events the state of the frequency process **Z**^*N*^ may change. Next, we define one by one how the different forces influence the sequence type frequencies.

### Virus Replication

At a virus replication event, two offspring are generated from a randomly selected virion. The selected virion itself vanishes and, to keep the population size constant, another random virus particle is chosen to be replaced by one of the offspring particles of the replicated virion. The offspring particles inherit the sequence type of the parent particle. Hence, if the sequence type of the offspring replacing a particle differs from the type of the replaced particle, we see jumps in the frequency process by 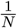 at the sequence type frequency of the replicating virus and by 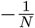 at the sequence type frequency of the replaced virus, both in the host in which the virus is replicating. For example, in Figure 1 (a) in the host, in which the virus replication event occurs, the frequencies of the sequence types 222 and 121 are 2/5 and 3/5, respectively, before the replication event. After the replication event, the frequency of type 222 has increased to 3/5 and the frequency of the sequence type 121 has decreased to 2/5. In the other hosts sequence type-frequencies remain unchanged. We assume that each virus particle is replicated at rate *γ*_*N*_ . For simplicity, we only assume binary replication. The theoretical results still remain valid as long as the offspring distributions are not too skewed, i.e., as long single virions do not give offspring to a macroscopic proportion of virions at a single replication event.

**Figure 1.**
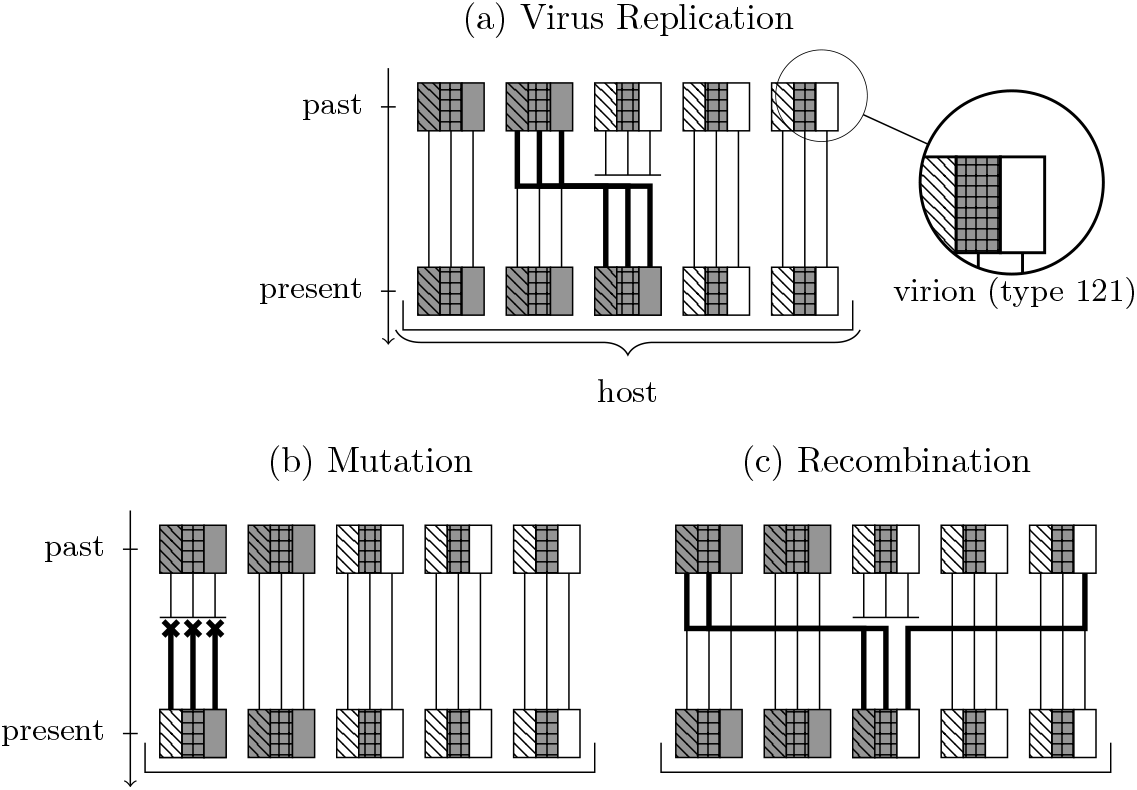
Illustration of the forces “virus replication”, “mutation” and “recombination” influencing the dynamics of the sequence type frequencies. Each host is infected with *N* virions and in each viral genome *L* loci are considered, which carry one of *K* genotypes. In this figure, *N* = 5, *L* = 3 and *K* = 2. Loci with a white background color carry genotype 1 and loci with a grey background color carry genotype 2.

### Mutations between genotypes

We consider parent independent mutation. Each virion is “hit” by a mutation at rate *µ*_*N*_ */N* . At a mutation event, the affected virus particle mutates to sequence type **k** = (*k*_1_, …, *k*_*L*_) for *k*_1_, …, *k*_*L*_ ∈ {1, …, *K*} with probability *p*_**k**_. Consequently, at a mutation event, the frequency of the sequence type **k** increases by 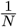 in the host affected by mutation, while the frequency of the sequence type **k**^*′*^, which the individual had before mutation, decreases by 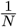. In Figure 1 (b) the frequency of the sequence type 222 in the affected host is 2/5 before the mutation event occurs. After the mutation event, the frequency of the sequence type 222 has decreased to 1/5 and the frequency of the newly appeared sequence type 122 is 1/5.

We do not require that mutations at different loci are independent. Instead, we only assume that, at a mutation event, the probability to mutate to a type *k* at any given locus is positive for any *k* ∈ {1, …, *K*} . More precisely, denote by

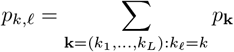

the probability of a mutation to type *k* at locus *ℓ*. We assume *p*_*k,ℓ*_ *>* 0 for any *k* ∈ {1, …, *K*} and any *ℓ* ∈ {1, …, *L*} .

Note that a mutation between genotypes considered here is not a mutation of a single base pair but rather the (much rarer) event that multiple base pairs mutate simultaneously. See the discussion section for more details.

### Recombination

Let **r** be a set of loci, i.e. **r** ⊆ [*L*]. At an **r**-recombination event in a host, a randomly chosen virus particle is removed and replaced by a recombinant that is generated by two viral particles that are chosen uniformly at random in the same host. The sequence type of the recombinant consists of the genotypes of the first viral particle at the loci of the set **r** and of the genotypes of the second viral particle at the loci of the set [*L*] \**r**. In Figure 1 (c) the frequencies of the sequence types 222 and 121 before the recombination event are 2/5 and 3/5, respectively. After the recombination event the frequencies of both, 222 and 121, are 2/5, and the newly appeared sequence type 221 of the recombinant has frequency 1/5. Each host is affected by an **r**-recombination event at rate *ρ*_*N*_ · *q*_**r**_. Let ℛ = {**r** ⊆ [*L*] : *q*_**r**_ *>* 0} be the set of recombination sets, for which a recombination event occurs wit h positive rate. We assume ∑_**r**∈ℛ_ *q*_**r**_ = 1. That is, the total rate at which a host is affected by recombination events is *ρ*_*N*_ .

### Reinfection

At a reinfection event, a viral particle in the reinfect*ing* host is replicated and the two offspring particles inherit the parent’s sequence type (and the replicated virion gets destroyed at replication). One of the offspring particles replaces a randomly chosen viral particle in a randomly chosen host (that we call reinfect*ed* host in the following). We see a frequency jump in the reinfected host by 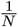 in the sequence type frequency of the reinfecting viral particle and by 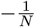 in the sequence type frequency of the replaced particle. For example, in Figure 2 (a) in the reinfected host (on the left hand side) the frequency of the sequence type 222 decreases from 1 to 4/5 and the frequency of the sequence type 121 increases from 0 to 1/5. In the reinfecting host, sequence type-frequencies stay (as in the other remaining hosts) constant. Each viral particle reinfects at rate *λ*_*N*_ */N* and hence each host reinfects other hosts at rate *λ*_*N*_ .

**Figure 2.**
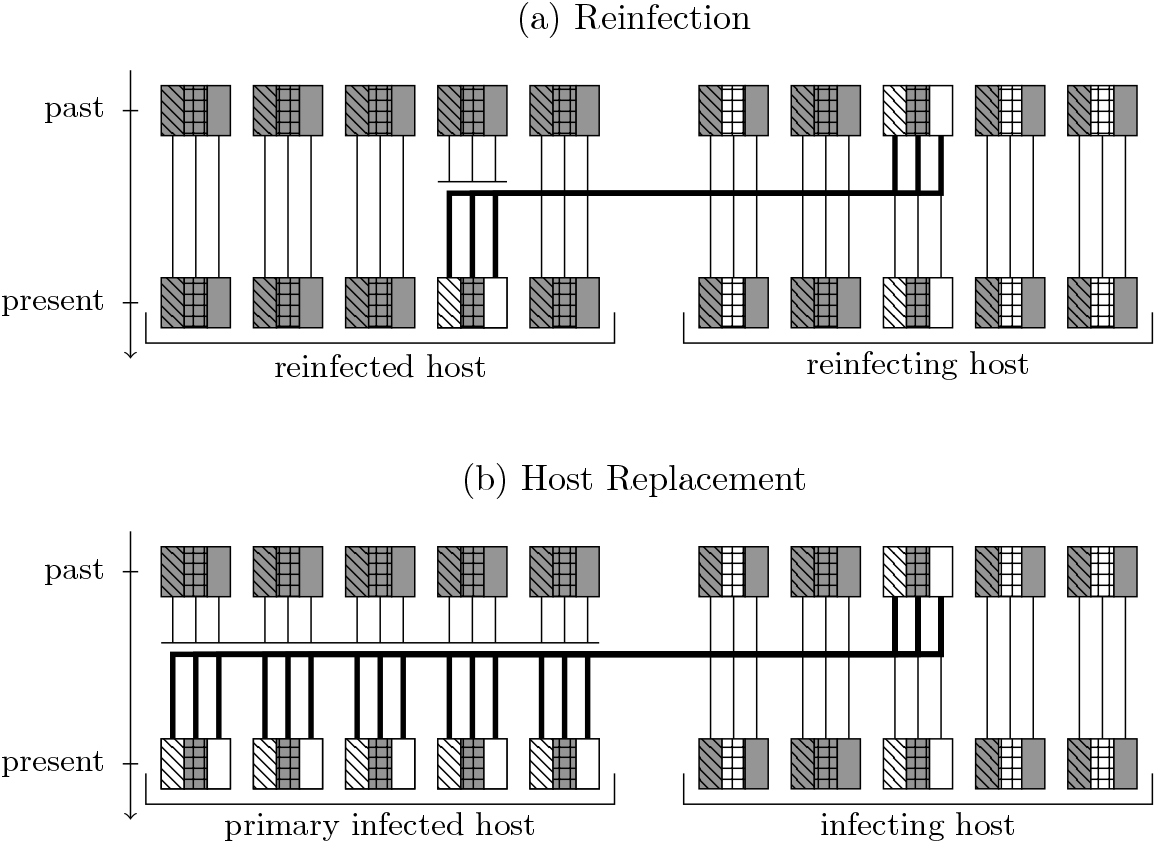
Illustration of the forces “reinfection” and “host replacement” influencing the dynamics of the sequence type frequencies. Each host is infected with *N* virions and in each viral genome *L* loci are considered, which carry one of *K* genotypes. In this figure, *N* = 5, *L* = 3 and *K* = 2. Genotypes with a white background color we call type 1 in text and genotypes with a grey background color we call type 2.

### Host replacement

At a host replacement event, a host dies and is replaced by a so far uninfected host that is instantly primary infected. At primary infection, a randomly chosen viral particle infects a so far uninfected host and during this event *N* viral particles are generated which are of the same type as the infecting virion. These *N* offspring particles found the viral population in the primary infected host. In particular, instantly after primary infection the host is infected with a single virus sequence type only. In Figure 2 (b) the host on the left hand side dies and is replaced by a primary infected hosts that gets instantly infected by the host on the right hand side. Before the event, the frequency of 222 in the host was one. The host is replaced and the primary infected host carries only type 121, i.e. the frequency of 121 in that host is one. The frequencies in the other hosts do not change. Host replacement events occur at rate 1 per host.

To analyze observed HCMV sequence data in the context of our mathematical model, we are interested in the sequence type frequencies in a sample of virions taken from a randomly chosen host. In [10], we have identified this asymptotic sequence type frequency spectrum as *N* → ∞ under the following assumptions: First, we make an assumption on the set of recombination events that happen at a positive rate. This assumption yields that linkage between every pair of loci can be broken by recombination with positive probability.

### Assumption R

For any *ℓ*_1_≠ *ℓ*_2_ ∈ [*L*] there is a set **r** ⊂ [*L*] with *q*_**r**_ *>* 0 such that either *ℓ*_1_ ∈ **r** *and ℓ*_2_ ∉ **r** or *ℓ*_1_ ∉ **r** *and ℓ*_2_ ∈ **r**.

Assumption **R** is fulfilled, for example, by the set of single-crossover recombination events, i.e. if *q*_{1,…,*ℓ*}_ *>* 0 for all *ℓ* ∈ [*L*]. Further examples are given in [10]. Furthermore, we make the following assumptions on the rates:

### Assumption A

1. The viral replication rate and the reinfection rate per host are of the same order. That is,

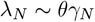

for some fixed *θ >* 0.
2. Reinfection and viral replication events occur on a faster time scale than mutation and recombination events as well as host replacement events. That is,

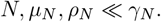
3. Recombination is faster than mutation:

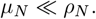

This assumption guarantees that with respect to genotype diversity the genealogies of two loci are asymptotically independent.
4. Mutation is strong enough:

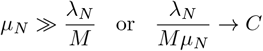

for some fixed *C >* 0. This assumption yields that loci are asymptotically *K*-allelic.

We discuss the applicability of these assumptions to the HCMV population in SI Text Section 1. In [10], sequence type frequencies under stationarity were identified for large viral and host population sizes. More precisely, the following statements are shown.

### Locus-wise type frequencies in the whole population

We define the locus-wise type frequencies in the whole virus population at sampling time by

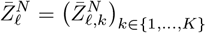

for *ℓ* ∈ [*L*] where 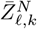 is the frequency of type *k* at locus *ℓ* in the whole population at sampling time, that is

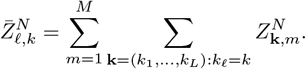

When Assumptions **R** and **A** hold, then (in the limit *N* → ∞) the locus-wise type frequencies 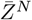 under stationarity can be described as follows: if 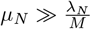, then 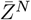 is asymptotically deterministic. More precisely, 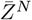 converges to a constant 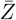 which depends on the parameters *θ* And 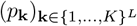. If 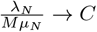, then 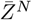 converges to a vector of *L* independent Dirichlet distributions. The parameters of the Dirichlet distributions depend on *θ, C* and 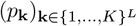, see Theorem 2.9 in [10] for details.

### Sequence-type frequencies in a randomly chosen host

We define the sequence type frequencies observed in a randomly chosen host

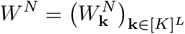

by 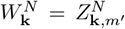 where *m*^*′*^ is a host chosen at random from {1, …, *M* }. Under Assumptions **R** and **A**, the distribution of *W* = lim_*N*→∞_ *W*^*N*^ is given by a Dirichlet distribution with parameter

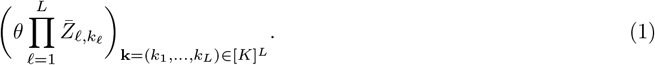

Here, 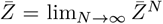. *W*^*N*^ can be interpreted as the sequence-type frequencies observed when sampling all *N* virus paricles from a randomly chosen host. Theorem 2.11 in [10] and the subsequent Remarks imply that not only the asymptotic distribution of *W* ^*N*^ but also the one that arises when sampling less than *N* (but still asymptotically infinitely many) particles from a random host follow this Dirichlet distribution.

### Parameter Estimation

Finally, Theorem 2.13 in [10] yields that the parameter in Equation (1) can be estimated by determining the empirical locus-wise type frequencies obtained when taking large samples of virions from a large number of hosts.

Together, the results suggests to fit a Dirichlet distribution to empirically measured genotype distributions. In the next sections we describe how we fitted the model to genotype data from HCMV.

## 2 Fitting the model to genotype data from the human cytomegalovirus

### 2.1 Modeling HCMV evolution

As assumed for the viral population in our model, HCMV is a herpesvirus that *persists* in its host after primary infection as well as after reinfection. That is, once infected, a host carries a HCMV population (established at possibly several infection events) until the end of its life. During persistence, the viral population switches between phases of reactivation and latency. In this manner, the virus is rarely detected by the host’s immune system in phases of latency but can spread at phases of reactivation. Complex mechanisms control these switches [11]. For simplicity, in the model presented in the previous section it is assumed that the virus is continuously replicating, that is we ignore phases of latency because in these phases the viral population does not evolve further. We nevertheless account for phases of latency by relating the viral replication rate and host replacement rate appropriately. For this purpose, we estimate (based on parameters available in the literature) the number of replication cycles happening within the lifespan of a typical host in SI Text Section 1.

In the host-pathogen model considered here, the genetic composition at a few genomic loci is analyzed. At all of these loci, pathogens are assumed to have one of a finite number of different types. For HCMV such a clustering into a few phylogenetically and genomically distinct genotypes has been observed for many DNA regions [2]. Some of these loci are known to encode proteins involved in cell-to-cell viral transmission and host cell entry, in strategies of immune evasion and immune modulation^1^, or in determining HCMV cell tropism^2^. Examples for this are UL73 and UL75 (encoding glycoproteins gH and gN), where different genotypes evoke different immune responses of the host [12, 13, 14], and UL55 (encoding glycoprotein gB), where they alter fusogenicity [15]. As in the model considered here, hosts have been shown to have mixed infections with different genotypes, e.g. at the ORFs UL6 [16], UL9 [17], UL55 [18, 19, 20, 17], UL73 [16], UL75 [6], UL139 [16], UL144 [17] and UL146 [17, 16].

For the analytical results from [10], recalled in the previous section, to hold, the Assumptions **R** and **A** on the model parameters need to be fulfilled. In SI text Section 1 we discuss which parameter regimes could be relevant for HCMV and if these regimes are in line with Assumptions **R** and **A**.

### 2.2 Data acquisition and description of the dataset

#### 2.2.1 Sample collection

Stored residual sample material from HCMV-DNA positive samples sent to the Center for Virology, Medical University of Vienna, for routine diagnostic testing between 2015 and 2022 (Ethical Approval No. 2156*/*2019) was collected fulfilling the following criteria: HCMV DNA load of at least 5 *×* 10^3^ copies/ml to ensure minor genotype/variant detection down to 5%, no known or suspected solid organ or stem cell transplantation. Each sample represents a unique individual.

Samples were selected randomly while taking age structure into account in order to obtain a sample representative of the persistently infected hosts with an active infection. It has been shown that older individuals are on average more often infected with several genotypes than younger individuals [6], which can also be observed in our dataset. This is reasonable because, on average, older hosts have presumably been reinfected more often than younger hosts. Since in our study the genotype distribution within hosts is of central interest, it is important to account for this age structure. Therefore we estimated the age structure of a representative sample, see SI Text Section 2 for details. This stands in contrast to many HCMV sample collections in the literature, where samples often stem from newborns, toddlers or transplantation patients and thus are not representative in terms of age distribution.

Clinical information about the patients was retrieved from medical records saved with the stored samples. Clinical signs or symptoms for diagnosis of HCMV by in-house quantitative PCR [21] are: suspected HCMV infection (*n* = 11), fever (*n* = 6), hepatopathy (*n* = 5), leuco-, thrombo-, neutropenia (*n* = 6), gastroenteritis with or without morbus crohn (*n* = 6), pneumonitis (*n* = 5), suspected malignant disease (*n* = 5), unknown (*n* = 20). Specimen types are plasma/serum (*n* = 50), urine (*n* = 8), respiratory secretions/lavage (*n* = 6).

#### 2.2.2 Selection of genomic loci and identification of signature amino acid residues within the coding sequences (CDS) UL4, UL75, UL78, US27

A search for genotypes was systematically performed at 153 different ORFs on the basis of the 101 whole genome sequences published in [1]. Annotations published together with the sequences were used to build phylogenetic trees for each ORF (based on the protein sequences, using MAFFT for alignment and SeaView for the generation of a phylogenetic tree). The results were filtered for trees and corresponding protein sequences which clustered into two clades. This was the case for the ORFs UL4, UL75, UL78 and US27, see Figures S1 to S12 in the Supplementary Material.

Dichotomous residues were re-evaluated in the dataset of 310 whole HCMV genome sequences that were publicly available at that time (09/2022). The following regions within UL4 (CDS: 46 - 149), UL75 (CDS: 1 - 87), UL78 (CDS: 5 - 91), US27 (CDS: 216 - 302) were finally selected. Reference nucleotide sequences are provided in the Supplementary Excel Sheet. Amino acid sequences from the sampled patients marked with the signature positions for genotype discrimination are provided in Figures S1 to S4.

#### 2.2.3 Primer design for UL4, UL75, UL78, US27 amplicon deep sequencing

*In silico* primer design was performed using Primer-Blast. Primers are located in highly conserved region up- and downstream of the selected regions by screening of a total of 310 whole genome sequences. Primers are listed in the Supplementary Table S3.

#### 2.2.4 Amplicon deep sequencing

DNA was isolated from patient samples using the EasyMag Nuclisense method according to the manufacturers’ protocol (BioMérieux). Briefly, 200 µl of patient sample was included into 2 ml of lysis buffer, and extracted with 50 µl elution buffer. For PCR, 20 µl mastermix containing 12.5 µl AmpliTaq Gold 360 Master Mix (#N8080241-Thermo Fisher Scientific), 0.5 µl primer each, and 6.5 µl nuclease-free water was combined with 5 µl of eluted DNA. Initial denaturation was performed at 95°C for 10 min followed by 40 cycles of 95°C for 30 s, annealing for 1 min at 49-51°C, and elongation at 72°C for 1 min, with a final extension time of 5 min at 72°C. Small aliquots of the PCR products were visualized on analytical agarose gels. Positive amplicons per samples were pooled equimolarly, then quantified by Qubit and subjected to library preparation and sequencing as described previously [22]. Briefly, 2 ng of pooled amplicon DNA was used to generate a 4 nM library using the Nextera XT library preparation and Index kit, followed by paired-end sequencing (2 *×* 150 cycles) on an Illumina MiSeq with automatic adapter trimming (Illumina, San Diego, CA, United States). FASTQ raw reads that passed filters were imported as paired-end reads into CLC Genomics Workbench 21.0 (Qiagen), trimmed by Q30 quality, and filtered for reads *>* 80 bp in length. After human genomic read removal, remaining reads were mapped to a set of about 10 reference sequences per considered ORF which can be found in the Supplementary Excel file with the default mapping parameters for match/mismatch scores, insertion/deletion costs, and a length fraction of 0.5 and a similarity fraction of 0.9. Consensus sequences of the mappings were extracted with a noise threshold of 0.05 and a minimum nucleotide count of 10 to insert ambiguity codes for single nucleotide polymorphisms. The resulting genotype consensus sequences were aligned, screened visually and unique sequences were selected for remapping. Genotype frequencies were determined by the number of reads mapped to the corresponding genotype sequences, expressed as a percentage of the total number of mapped reads. Amino acid alignments of unique sequences are provided in Figures S1 to S4.

### 2.3 Methods and data analysis

#### 2.3.1 Preparation of the dataset

For the analysis we used CMV material contained in serum, urine, EDTA-Plasma, respiratory secretions and lavage sampled from *n* = 64 different individuals, sent to the Center for Virology, Medical University of Vienna, for routine diagnostic testing. Genotype distributions at the four open reading frames UL4, UL75, UL78 and US27 within the samples were determined, see Section 2.2.4 for details. According to a phylogenetic analysis applied to the 101 whole genome sequences provided with [1], at each of these four loci the CMV sequences cluster into two genotypes, as described in Section 2.2.2. For the following, we define a sequence as type 1 at a given locus (e.g. at locus UL4) if the amino acid sequence at that locus is of the same type as that of the Merlin strain^3^, and as type 2 otherwise. A piece of viral DNA sequenced at *L* loci can thus be represented as a length *L* string of 1’s and 2’s. For *L* = 4 and the loci UL4, UL75, UL78 and US27, the Merlin sequence is then, for example, of type 1111, while the AD169 reference sequence^4^ is of type 1212. We denote the set of all such sequence types (of viral particles) by *S* = {1, 2} ^*L*^ (= {1111, 1112, …, 2222}, when *L* = 4) and the frequencies at which these sequence types are present in sample *i* by

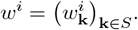

Using the mapped reads per genotype sequence we can quantify the amount of a genotype at single loci in a sample but due to the short read length, we lack linkage information across loci and therefore the frequencies *w*^*i*^ are not available. In the following we denote the (available) frequency of genotype 1 at locus *ℓ* ∈ {1, …, *L*} in the material sampled from individual *i* by 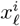. In particular,

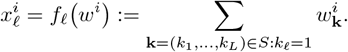

All measured frequencies can be found in Supplementary Table S4. From these frequencies, we estimate the genotype frequencies at each locus in the total population of CMV infected individuals by taking the mean frequency for each locus. We denote the estimated frequency of genotype 1 at locus *ℓ* by 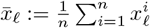. The resulting frequencies for loci UL4, UL75, UL78 and US27 are summarized in Table 1.

**Table 1.**
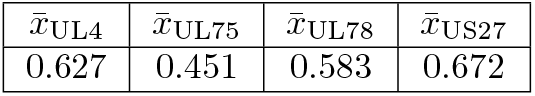
Estimated Genotype Frequencies in the Total Population.

Note that the sequence type frequencies *w*^*i*^ cannot be inferred from the measurements of 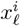 in general due to missing linkage information. For example, it is unclear which types of sequences are contained in sample 64 of Supplementary Table S4. All sequence types except of those with type 2 at the third locus are possible.

Further research is necessary to evaluate how well within-host type frequencies are estimated by the measured frequencies in the samples. CMV sequences can only be detected in samples if the viral population size is sufficiently large. This is the case when the virus is being replicated intensively. In the persistently infected adult host, this occurs at reactivation and reinfection events. Frequencies could be biased to genotypes of viral particles that initiated the reactivation/reinfection event. Furthermore, only one type of material (e.g. urine) was collected per individual and the genotype frequencies in that material may not accurately represent the frequencies in this host’s whole viral population.

Since the measurements of the frequencies can be rough and biased at the boundaries 0 and 1, we divide genotype frequencies into three groups, which we call “infection patterns” in the following. For a cutoff-value *c* ∈ (0, 0.5), we define that, at a locus in a sample, *mainly type* 1 *is present* if the frequency of type 1 is at least 1 − *c, both types are present* if the frequency of type 1 lies between *c* and 1 − *c*, and otherwise *mainly type* 2 *is present*. Via maximum likelihood estimation, we determine a cutoff-value *c* and the model parameter *θ* under which the data set is most likely, see Subsection 2.5.2 for more details. The different infection patterns are abbreviated by 1, *m*, and 2, where *m* stands for mixed. More formally, we set

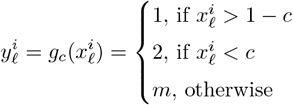

Applying this transformation to the frequencies from Supplementary Table S4 with (the according to the maximum likelihood estimation best fit value) *c* = 0.07 results in Supplementary Table S5.

### 2.4 From sequence type frequencies to infection patterns

Recall from Section 1 that, under appropriate rate assumptions, the sequence type frequencies *W*^*N*^ in a sample taken from a randomly chosen host asymptotically follow a Dirichlet distribution depending on the single locus type frequencies 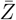 in the whole population and the parameter *θ*. We estimate 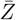 by 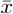 and infer the parameter *θ* (together with the cutoff-value *c*) by maximum likelihood estimation as described in Section 2.5.2. *W* is a random variable taking values in Δ = {*w* = (*w*_**k**_)_**k**∈*S*_ ∈ [0, 1]^*S*^ : ∑_**k**∈*S*_ *w*_**k**_ = 1 }. We apply the functions *f*_*ℓ*_ defined above to *W* to arrive at a vector of genotype frequencies (for the *L* loci) *X* = *(f*_1_(*W*), …, *f*_*L*_(*W*)) ∈ [0, 1]^*L*^.

For classifying *W* into a vector of infection patterns 1, *m* and 2 given a cutoff-value *c*, we proceed similarly as above by applying the function *g*_*c*_ on each component of *X*. The full infection pattern (for all loci) depending

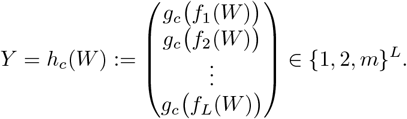

See also Figure 3 for an overview of the transformations we applied to the model-derived distribution and to the dataset. *Y* is a discrete random variable whose distribution depends on 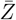, *θ* and *c*. More precisely, the probability weights 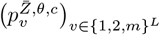 of the distribution of *Y* are given by

**Figure 3.**
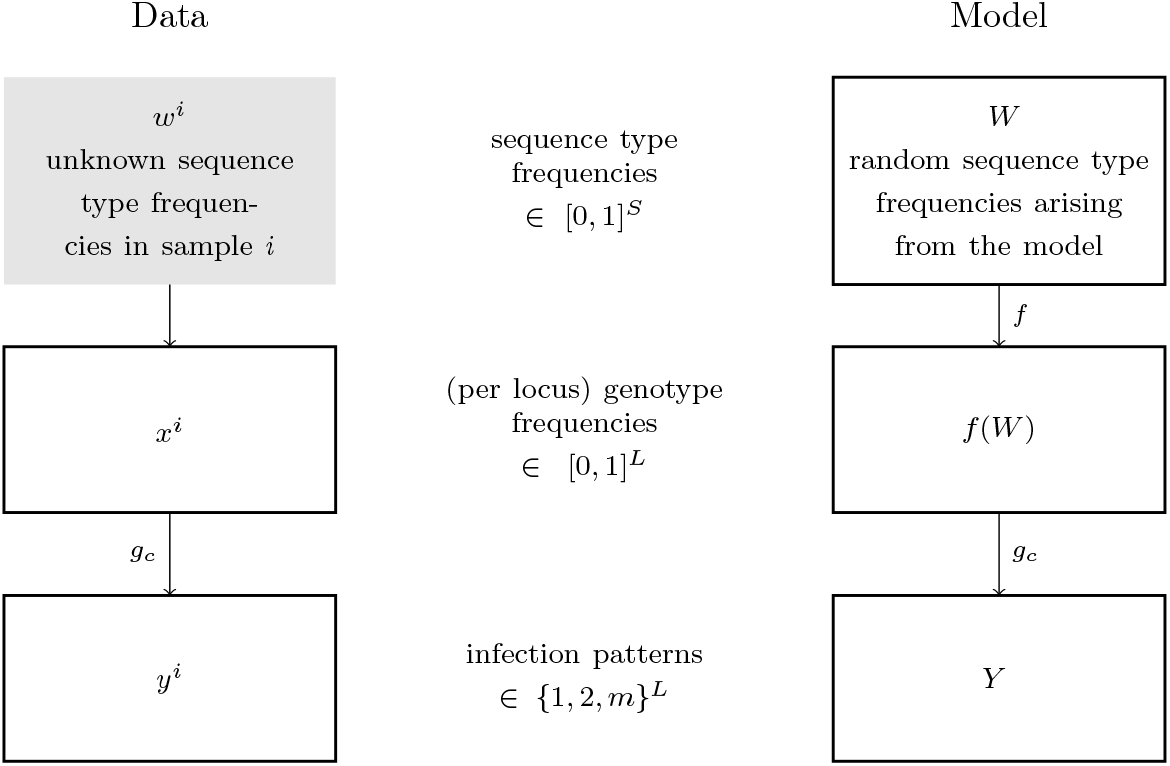
Overview of the transformations applied to the dataset and to the distribution arising from the model.

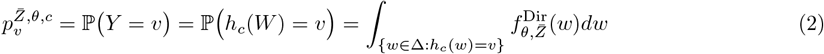

where 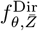 is the density of a Dirichlet distribution with parameter 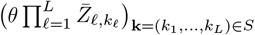.

### 2.5 Inference

Since the number of sampled hosts is small in comparison to the total number of hosts, we assume that our (transformed) data points *y*_*i*_ are independent draws from the same discrete distribution on {1, 2, *m*}^*L*^, which is given by a vector of probability weights 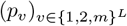. Note that the actual, unknown probability vector 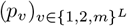 may or may not arise from our model, since the set of all probability vectors 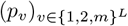 is higher dimensional than the set of allowed vectors resulting from our model. We will use maximum likelihood estimation to determine a model parameter 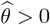 and a cutoff-value 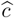 which best describe the observed data.

#### 2.5.1 Monte-Carlo Simulation

The goal is to identify a best fit parameter combination 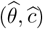 (given 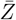). Given an arbitrary parameter combination 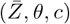 it is, however, analytically not straightforward to compute the corresponding vector of probability weights 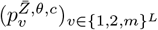. We therefore determine the weights numerically using Monte Carlo simulation. More precisely, for a fixed choice of *θ, c* and 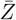 we simulate *d* ∈ ℕ i.i.d. realizations of *W*, named 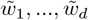 and apply the function *h*_*c*_ to each realization. If *d* is large enough, then, by the law of large numbers,

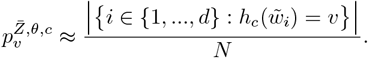

#### 2.5.2 Maximum-Likelihood Inference

We assume that the data (i.e., the rows of Table S5) arise as independent draws from the same unknown distribution on {1, 2, *m*}^*L*^. Under this assumption, the corresponding value counts given in Table S6 are multinomially distributed according some probability vector 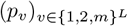.

Using maximum likelihood estimation, we find values for *θ* and *c* (given 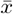) that best match the observation, i.e. we determine

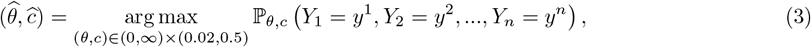

where *Y*_1_, *Y*_2_, …, *Y*_*n*_ are i.i.d. according to the probability vector 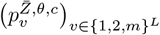 . We consider only cutoff-values *c* of at least 0.02 because the detection threshold for the qPCR-method used for sequencing the HCMV samples is 1 *×* 10^2^ copies/ml for each genotype [23], and in each sample at least 5 *×* 10^3^ viral DNA copies/ml were detected.

### 2.6 Goodness of fit

Next to a visual comparison, we assess the goodness of fit by computing *p* values as the overall goal is to decide whether the proposed model may be a reasonable null model. A high *p* value has the interpretation that the observed data are a “typical” observation in the inferred model and supports the belief that the proposed model accurately describes the viral dynamics underlying the observation.

We, again, use Monte-Carlo simulations to determine the total probability to observe a realization that has equal or smaller probability than the probability of the observation, assuming that the data follow the inferred distribution. This probability is

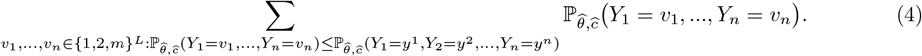

Due to symmetry (the random variables (*Y*_*i*_)_*i*=1,…,*n*_ are i.i.d.), this is equivalent to computing *p*-values for a multinomial distribution. In our case this multinomial has, however, |{1, 2, *m*} ^*L*^ = 3^4^ = 81 categories and 48 trials (see Section 2.7.1), rendering the computation of exact *p*-values infeasible due to the high dimensionality of the outcome space. Instead, we simulated a large number *d*^*′*^ ∈ ℕ of i.i.d. samples from a multinomially distributed random variable *R* with number of trials *n* and probability weights 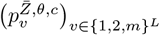. By the law of large numbers (4) can be approximated by

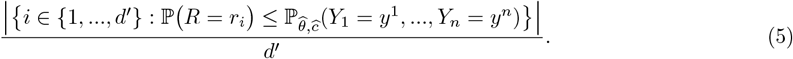

### 2.7 Results of the Inference

#### 2.7.1 Inference on all four loci

We first carried out the maximum likelihood inference, as described above, using data from all four available loci. From the 64 samples that were initially in the dataset, we only considered the 48 samples for which genotype frequencies at all four loci were available. Using the remaining data points, we calculate the likelihoods on the right-hand side of Equation (3) depending on *θ* and the cutoff parameter *c* for *θ* = 0.01, 0.012, …, 0.21 and *c* = 0.005, 0.01, …, 0.1. The resulting approximations of the likelihoods are illustrated in Figure 4 (a). As estimators for *θ* and *c* we use the maximizing values, which were 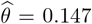 and 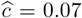 and are marked by a red cross in Figure 4 (a). The corresponding probability weights 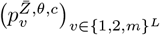 on the infection patterns were approximated using Monte-Carlo simulation as described in Section 2.5.1.

**Figure 4.**
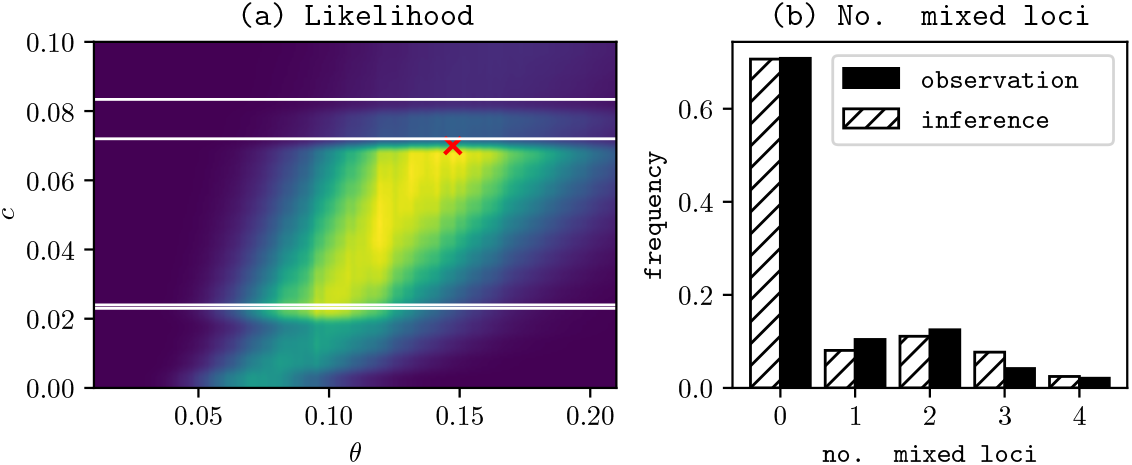
(a) Monte-Carlo approximation of the likelihood function depending on the model parameter *θ* and on the cutoff parameter *c*, with maximum likelihood estimates 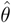 and 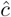 marked by a red cross. Note that the likelihood seems to depend continuously on *c* but not continuously on the cutoff parameter. The discontinuities in the cutoff parameter occur at those levels, at which the changed cutoff leads to a change in the classification of the raw genotype frequencies (indicated by white lines). (b) Plot of the number of mixed loci observed in the dataset and predicted by the inferred model.

In Figure 5, these probability weights are plotted together with the empirical probability weights observed in the data. There are 3^4^ = 81 possible infection patterns but only 48 data points, so we can not hope for a perfect fit of the data with the model. However, in Figure 5, many of the most frequently observed infection patterns also showed a high theoretical probability weight. To further asses the fit, we again used Monte-Carlo simulation as described in 2.6 to obtain an approximated *p*-value of 0.38322. Additionally, we plotted the inferred distribution on the number of mixed loci against the numbers of mixed loci from the sample in Figure 4 (b).

**Figure 5.**
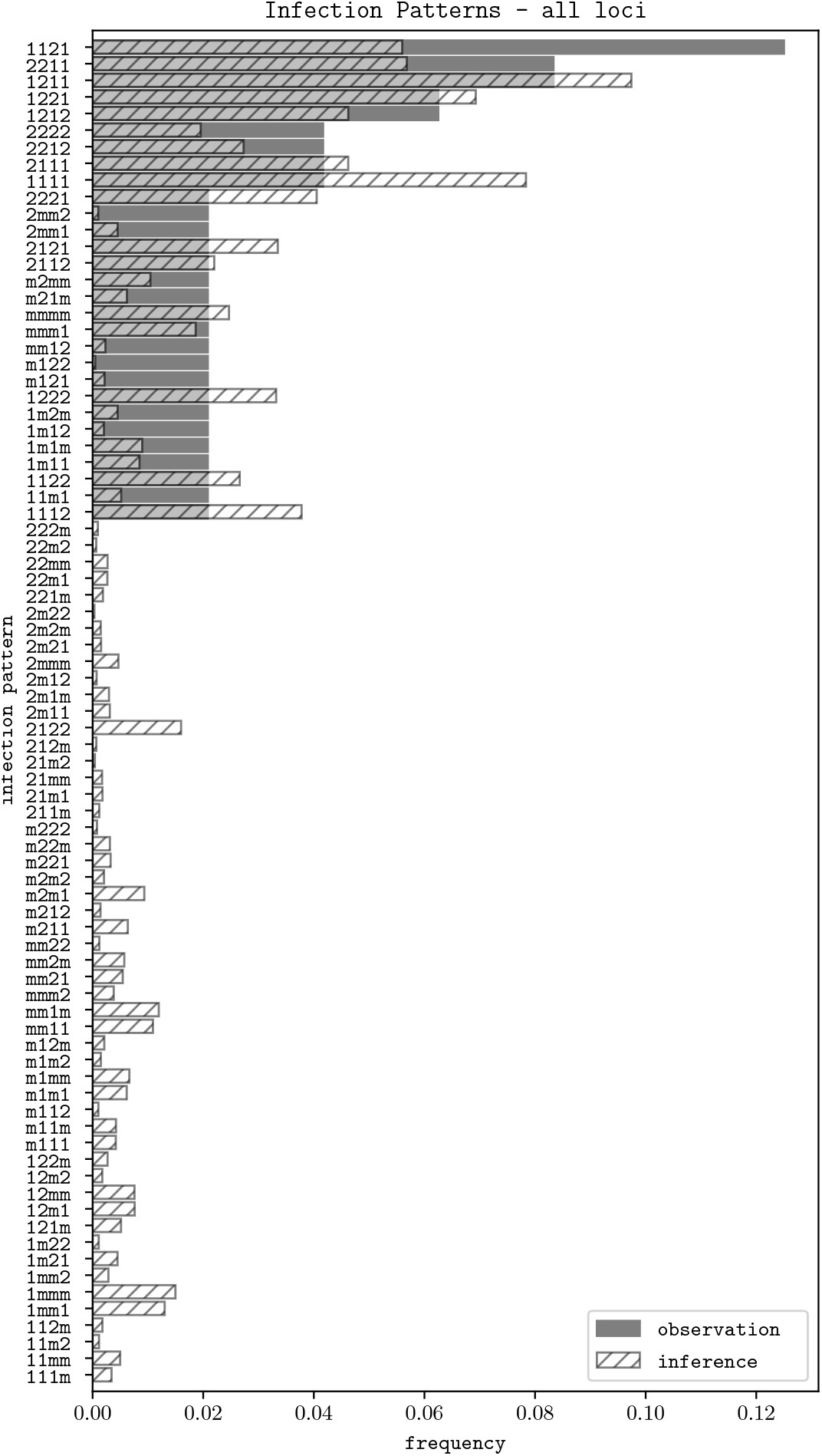
Plot of the empirical probabilities of each infection pattern (“observation”) with the probability weights in the fitted model (“inference”).

#### 2.7.2 Consistency with fewer loci

Above we have inferred the best fit model parameters 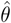 and 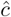 using data, for which genotype frequencies were available at all four loci. The parameter 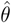 can be interpreted as the ratio of the reinfection and the viral replication rate, and 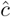 can be interpreted as cutoff parameter or detection threshold. If the proposed model fits well, then both parameters should not depend on the studied loci. To check this, we also fitted the model to only a single locus. The estimated values of *θ* and *c* for the different loci can be found in Table 2. Except for locus UL78 the estimated values of *θ* and *c* lie in the same range as when all loci are used for fitting. Thus, using an arbitrary subset of the four loci UL4, UL75, UL78 and US27 and comparing the observed infection patterns with the model prediction with the *same* parameters 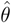 and 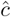, we should observe a good fit. Se some data points are missing due to technical issues (no amplicon or no sequencing reads), see Supplementary Table S4, more samples become available when considering fewer loci. These additional samples have not been used for maximum likelihood estimation of 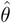 and 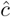 and thus provide new information.

**Table 2.**
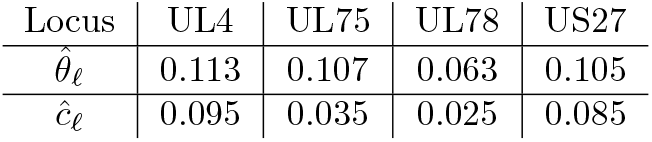
Best fit parameters estimated using information from single loci.

| Locus | UL4 | UL75 | UL78 | US27 |
| --- | --- | --- | --- | --- |
| $\hat{\theta}_\ell$ | 0.113 | 0.107 | 0.063 | 0.105 |
| $\hat{c}_\ell$ | 0.095 | 0.035 | 0.025 | 0.085 |

In the following, we study all subsets of the four loci and compare the fitted model with the observed infection patterns visually and with approximated *p*-values.

##### Three Loci

First, we study all subsets containing exactly three loci. The corresponding *p*-values are approximated using the same method as in Equation 5 and are given in Table 3.

**Table 3.**
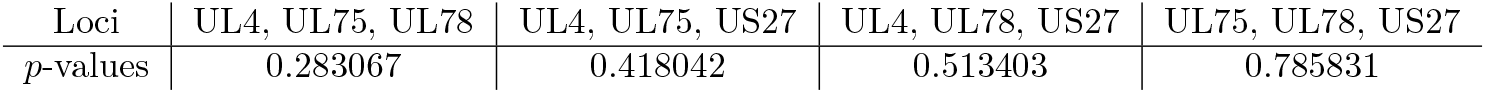
*p*-values observed when the model with the inferred parameters 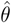 and 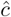 is applied to any three loci. Detailed plots for the combination of loci with the largest *p*-values are given in Figures 6 (a) and 7 (a). Plots for the loci with the lowest *p*-values are given in Figures 6 (b) and 7 (b).

In Figure 7, we plotted, similar to Figure 5, the observed and inferred infection patterns for the combination of loci that achieved the highest and the lowest *p*-values in Table 3, respectively. The best and the worst fitting combination of loci both show a surprisingly good fit.

In order to summarize the diversity observed at three loci, we also plotted the proportion of individuals carrying both genotypes at 0, 1, 2 or 3 loci in the dataset and in the fitted model in Figure 6.

**Figure 6.**
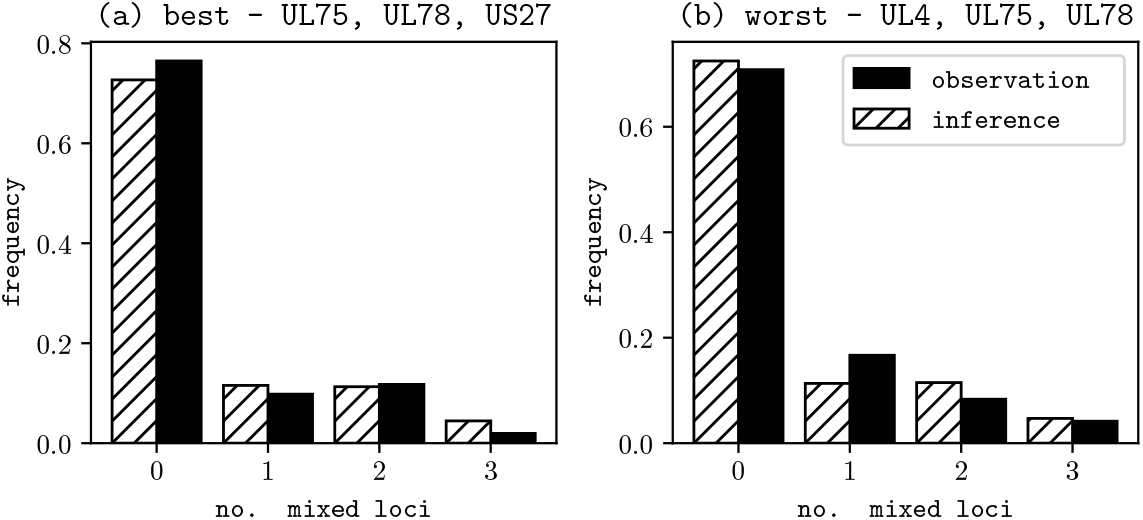
Plot of the number of mixed loci observed in the dataset and the ones predicted by the inferred model when three loci are used for fitting. (a) Combination of loci where the best fit is obtained. (b) Combination of loci with the worst fit.

##### Two Loci

For combinations of two loci, we find, on average, even higher *p*-values, see Table 4. Again, we compare the inferred model with the data visually via infection patterns and via the numbers of loci with both genotypes presentin Figure 8.

**Table 4.** *p*-values observed when the model with the inferred parameters is applied to any two loci. Below, we illustrate the combination of loci with the lowest and highest *p*-values in Figure 8

| Loci | UL4, UL75 | UL4, UL78 | UL4, US27 | UL75, UL78 | UL75, US27 | UL78, US27 |
| --- | --- | --- | --- | --- | --- | --- |
| $p$-value | 0.852244 | 0.682135 | 0.823335 | 0.843856 | 0.382849 | 0.936797 |

**Figure 7.**
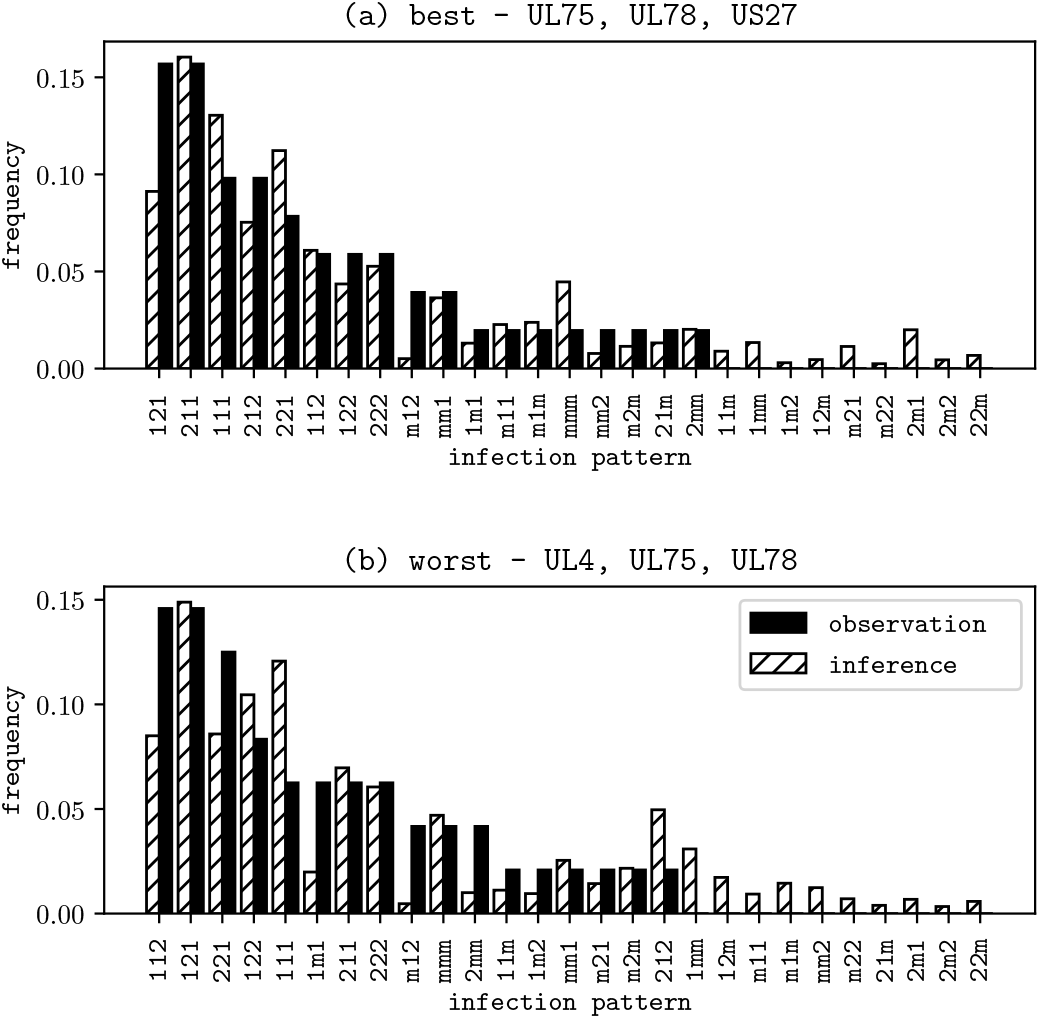
Empirical frequencies and theoretical probabilities of the infection patterns for the best(a) and worst(b) fitting combination of three loci.

**Figure 8.**
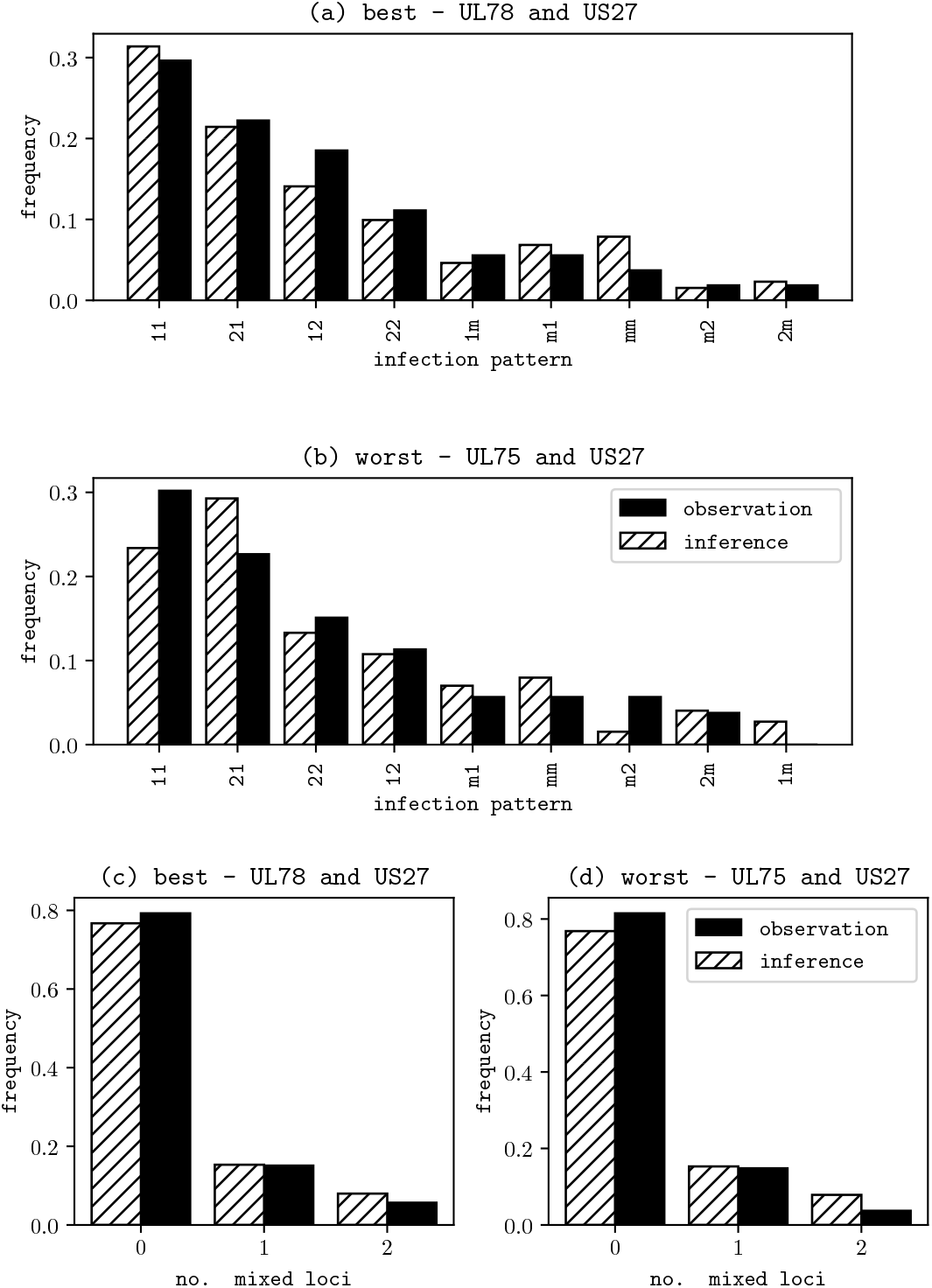
Top: empirical frequencies and theoretical probabilities of the infection patterns for the best (a) and worst (b) fitting combination of two loci. Bottom: the numbers of mixed loci observed in the dataset and the ones predicted by the inferred model when two loci are used for fitting. On the left hand side (c) the combination of loci is shown where the best fit is obtained and on the right (d) the combination of loci with the worst fit.

##### Single Loci

We give the *p*-values observed for single loci in Table 5 and observe a similarly high average *p*-value as in the case of two loci.

**Table 5.** *p*-values observed when the model with the inferred parameters is applied to single loci.

| Locus | UL4 | UL75 | UL78 | US27 |
| --- | --- | --- | --- | --- |
| $p$-value | 0.866256 | 0.983763 | 0.516066 | 0.761349 |

When focusing only on one locus, information on the number of mixed loci is directly available from infection patterns, hence we only plot inferred and observed infection pattern distributions in Figure 9.

**Figure 9.**
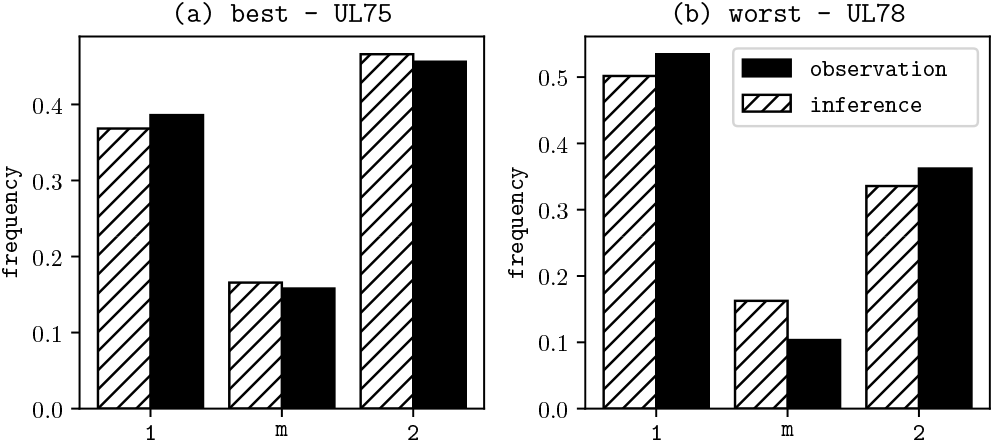
Best and worst fitting infection patterns for single loci.

## 3 Discussion

Many host-pathogen pairs share a long-term evolutionary history. In many such systems, infections of healthy hosts are common and asymptomatic, at least in the short term. From an evolutionary point of view, it is reasonable to assume that these pairs reached a steady state and evolution is mainly neutral.

In [10], a model was developed for the neutral, long-term evolution of a virus population that is distributed across its host population and capable of persistence in its host as well as of reinfection. Under certain parameter assumptions on the viral replication, recombination and mutation rates as well as on the host and within host viral population sizes, see Assumptions **R** and **A** in Section 1, the asymptotic sequence type frequency spectrum for a set of genomic loci, at which sequences cluster into few types, has been determined. Building on these results, we here developed a method to fit this model to human cytomegalovirus DNA data.

Human cytomegalovirus is an ancient herpesvirus that co-evolved with its human host for millions of years [24]. Furthermore, it has been shown that DNA sequences at many genomic loci, more precisely open reading frames (ORF), cluster into several so-called genotypes [2, 3]. This suggests that the model from [10] might be appropriate for representing the evolution of human cytomegalovirus.

To apply the fitting method to genotype data from HCMV, we searched for ORFs which cluster mainly into two genotypes. This led us to consider the ORFs UL4, UL75, UL78 and US27. Based on the 101 whole genome sequences from [1], we determined for each locus sites at which the two genotypes can be identified uniquely. For some ORFs like UL75, it has been shown that the two genotypes encode for different epitopes. For many other loci, however, it is not clear, if the different genotypes create functionally different virions and even if the clustering into genotypes is just a result of genetic drift or if selection is forming this clustering.

Importantly, our method can be applied not only to loci with exactly two genotypes but also to loci with more variants. However, in this case, it is preferable to have a larger dataset as the number of possible infection patterns grows quickly with the number of genotypes.

Since age likely influences the sequence type distribution within a host, we used statistics available in the literature to estimate the age structure of humans shedding HCMV and took samples from these individuals (roughly) according to this distribution. This representative sampling is appropriate for studying the long term evolution of HCMV and stands in contrast to many HCMV studies, which often only consider specific groups of hosts, e.g. infants, pregnant women or transplant recipients.

The results in [10], which we use to fit the model to DNA data from HCMV, are based on the parameter Assumptions **A** and **R**. In the Supplementary Material, we assess the validity of these assumptions for HCMV in detail by comparing them with estimates found in the literature. Previous research clearly supports the validity of some of the assumptions. Exemplary for this are genome-wide recombination and mutation studies which yield evidence for recombination breakpoints between all four studied ORFs (i.e. the validity of Assumption **R**) as well as for recombination and mutation rates to be much smaller than viral replication rates (i.e. part of Assumption **A**, 2.). Other assumptions are harder to evaluate, for example part of Assumption **A** which states that the effective viral population size within host is much smaller than the number of viral generations in a host. In this context it is important to note that the model aims to represent the evolution of the *persistent* viral populations within the hosts. Albeit complicating parameter assessment, this approach captures the idea that virions generated at reactivation events probably only contribute little to the evolution within the host in which they are generated, but are mostly relevant for primary and re-infections, because the host’s immune response eliminates most virions.

Taking into account that only few parameters were fitted in comparison to the large number of possible infection patterns we find a (surprisingly) good fit of the model. None of the computed *p*-values turned out particularly low, indicating that the observed infection patterns are rather typical outcomes of the fitted model, see Section 2.7.1 and Tables 3 to 5. During the fitting procedure we infer the relative frequency of viral replication in comparison to reinfection. Our parameter inference yields that viral replication is roughly seven times more frequent than reinfection. This estimate should be validated with further data. At a first glance, the inferred reinfection rate seems to be rather high and it indeed exceeds previous estimates. As discussed in the SI Text Section 1, it is, however, not obvious how to measure the reinfection rate. Methods used in the literature seem to under(and also over-) estimate this rate. In particular, in contrast to methods used so far, it might be useful to consider larger genomic regions, since at reinfection events also diversity that is already present in the host can be transmitted. Methods to estimate reinfection rates based on genomic diversity might otherwise not detect a reinfection event.

Since the elderly population in general sheds virus more often than younger adults and since these hosts typically might harbor a relatively rich genomic diversity of HCMV, the elderly population could contribute significantly to reinfection events at which further genomic diversity of the HCMV population is transmitted to the reinfected host. Especially for vulnerable hosts, it could be important to figure out how frequent reinfection events are and which contacts are the most risky ones, since at reinfection events also variants with an altered biological behavior [25, 26] could be transmitted.

A characteristic feature of HCMV infections is the ability to switch between phases of latency and active replication. In the model considered here, phases of latency are only accounted for by scaling down the viralreplication rate. For simplicity, it is also assumed in the model that the size of the persistent population is constant. Further population genetic analysis is necessary to better understand the within host population dynamics of the persistent population. The difficulty in studying this population is that it (usually) cannot be sampled directly as it is mainly located in the myeloid lineage cells. Sampled virus often is collected at phases of active replication, e.g. in plasma, urine or saliva. Due to the exponential growth during these phases single viral strains could be overrepresented in these samples. Time sampled data could help to infer information about the persistent population, because different parts of the persistent population could initiate reactivation at different time points.

In the model from [10] at a locus a finite number of different genotypes are possible. This assumption represents the picture observed for HCMV sequences. For a couple of these loci, it has been clarified that different genotypes evoke different immune responses or the like. For many loci, however, it is not clear if the separation of genotypes is just a result of genetic drift, i.e. random neutral evolution, or if other variants are not viable.

In the latter case an alternative to the neutral scenario considered here that could explain the high observed diversity of HCMV would be that different genotypes are maintained in the viral population due to balancing selection. For example immune evasion strategies may lead to such an effect. Different antibody responses exist against the antigens of genotypes gH1 and gH2 of the glycoprotein gH (UL75) and against two different antigenes of gB (UL55) [12, 27]. Both genotypes may be controlled according to some predator-prey-dynamics caused by the immune system and due to a restricted amount of cells which can be latently/persistently infected. Further research is necessary to investigate if such dynamics shape HCMV evolution. From an applied point of view this could be of relevance because it would mean that diversity is actively maintained in the HCMV population. Also resistant genes against drugs could hitchhike in such a diverse population, even when the drug treatment is paused. In a neutrally evolving population diversity would drift out of the population more easily.

A first step into this direction has been previously achieved for a one-locus model with two genotypes [8]. In a multi-locus variant of this model, it would be reasonable to assume that the frequencies, at which the types at a locus are balanced, are locus-specific and that selection parameters can differ between loci. In this case, the model can be perfectly fitted to any sequence-type frequency data, at least if the recombination rate is sufficiently high (in contrast to our neutral model, where we fit only the two parameters *c* and *θ*). Hence, to investigate if HCMV evolution is not only neutral (as considered here), but also significantly impacted (at least at some ORF) by selective forces a finer analysis and more data are necessary. In particular, time-sampled data could be helpful.

## Footnotes

1 i.e., viral mechanisms to interfere with the immune response of the host

2 i.e., allowing HCMV to replicate in different cell types of the host

3 https://www.ncbi.nlm.nih.gov/nuccore/AY446894

4 https://www.ncbi.nlm.nih.gov/nuccore/X17403

