## Supplementary Material for "Analyzing Human Cytomegalovirus Genotype Diversity by Modelling Viral Population Dynamics"

**Supporting Information**

Raphael Eichhorn, Irene Görzer, Büsra Külekci, Madlen Mollik, Cornelia Pokalyuk

**Remark:** The code used for inference and goodness of fit can be found at `https://github.com/raeichho/hcmv_data_analysis`.

#### 1 Applicability of the model to HCMV

In this section we discuss the model design in the context of HCMV. In particular, we identify the parameter regimes in which the different evolutionary forces could be acting and interpret the best fit parameter estimate $\widehat{\theta}$.

Most studies in the literature focus on HCMV infections in transplant recipients and congenitally infected children, where viral replication is often more pronounced than in immunocompetent hosts. On an evolutionary time scale these patients are probably of low relevance, therefore here we primarily refer to the literature analyzing the infection dynamics in healthy hosts.

##### Viral population size within hosts

We assume in our model that the persistent viral population size within hosts is constant. With this assumption we implicitly assume that there exists some carrying capacity that the viral population maintains in the host also at phases of latency. For simplicity we assume that this carrying capacity does not change over time and does not differ between hosts. It is currently unknown if these model assumptions are reasonable for HCMV. The size of the persistent population may be increasing as long as the immune system cannot control HCMV sufficiently (the controllability of HCMV depends on the age of the host: in young, infected children with and without symptoms high HCMV DNA loads can be detected for more than a year [1], in adults the virus sporadically reactivates [2, 3]). It is possible that the viral population size within a host is decreasing over time when the virus does not reactivate or when the host is not reinfected sufficiently often, since cells in which the virus persists are replaced. The host immune response on HCMV certainly differs between hosts. This could also affect the carrying capacities. However, at least the number of latently infected mononuclear cells, which (in particular the subset of the pluripotent CD34+ bone marrow progenitor cells (BMC) [4]) are thought to be the main site of the persistent HCMV population, did not vary much in eight different, presumably immunocompetent hosts [5]. Albeit the number of examined patients is rather small, this finding can be taken as an indication for rather uniform carrying capacities among hosts.

Currently, no estimates of the effective size of the persistent viral population in the healthy host are available. Only the population histories of congenitally infected infants, in which a latent phase has not been reached yet, have been inferred. The effective compartmental population sizes of the HCMV populations were estimated to be of order 1000 [6].

We estimate the actual number of virions latently infecting a host to be similarly high, namely roughly between $7 \cdot 10^3$ and $2 \cdot 10^5$. We estimated the actual number of virions as follows: In [5], 0.004-0.01 % of granulocyte colony-stimulating factor (GCS-F)-mobilized BMCs[1] have been estimated to be infected with HCMV and in this study HCMV-infected cells contained on the order of 1 to 13 genomes per cell. In [7], 0.1±0.02% of CD 34+ BMCs have been estimated to be infected with HCMV in HCMV-seropositve healthy hosts. The number

[1] CD34+ BMCs are mobilized with GCS-F to migrate to the peripheral blood where they can be sampled more easily. There is no indication that the proportion of CD34+ BMCs infected with HCMV is altered by this treatment

of HCMV genomes per infected cell have not been estimated in this study. The majority of CD 34+ cells are contained in the bone marrow. The estimated mean number of CD 34+ BMCs per kg body weight is $2.4 \cdot 10^6$ [8]. Assuming a body weight of 70 kg for a standard individual, we estimate that the total count of HCMV virions contained in CD 34+ BMCs lies between $0.00004 \cdot 70 \cdot 2.4 \cdot 10^6 \approx 7 \cdot 10^3$ and $0.0001 \cdot 13 \cdot 70 \cdot 2.4 \cdot 10^6 \approx 2 \cdot 10^5$. These are estimates for physical size of the persistent population. The effective population size may be lower than that.

#### Population structure within hosts

Migration rates between compartments have been estimated to be small in HCMV samples from congenitally infected infants [6]. With our model we aim to represent the evolution of the persistent viral population which is thought to be located primarily in CD34+ bone marrow cells, see above. The bone marrow is distributed over several bones. This population structure could also translate to the persistent CMV population. Currently, no estimates of such a population structure are available so that the simplest assumption of panmixia should be preferred.

#### Population structure of the total viral population

So far only few studies considered the geographic structure of the HCMV population. In [9], 128 full genome sequences have been analyzed with 125 sequences from Europe, one from the USA, one from South Korea and one from China. The estimated phylogeny exhibits a star-like form, and the sequences outside of Europe did not cluster separately. The worldwide distribution for UL73 (for samples from Europe, China, Australia, and Northern America) was analyzed in [10], and for UL146 und UL 139 (for samples from Africa, Asia, Australia, and Europe) in [11]. No significant differences were detected in the genotype distributions. HCMV population structure on a genomic scale was studied recently in [12]. Evidence for a geographic structure of the HCMV population was found mainly in conserved genomic regions which are not the focus of this study. In particular, three out of the four loci used for our analysis were classified as "nongeographic". Therefore the simplistic assumption of a panmictic, i.e. well-mixed, host population might be realistic.

#### Size of the host population

The effective population size of the human population is estimated to be of order $10^3$ to $10^4$ [13, 14]. However, the effective host population size relevant for HCMV evolution could be higher because the genealogy of the viral population is not only influenced by host reproduction events, but also by reinfection events leading to a higher level of admixture (of within host populations). This should be reflected in a higher effective host population size. The size of the host population is naturally upper bounded by the current size of the human population which lies at around $8 \times 10^9$.

#### Viral replication

We model the evolution of a replicating *persistent* viral population. In reality, the replication rate obviously is not constant. There are phases at which the immune system essentially silences viral replication and phases at which the virus reactivates. In phases of reactivation, virus is produced and shed into bodily fluids to infect other hosts, but also cells are infected in which the virus eventually will persist. In our model, the time for each host is tacitly rescaled to the speed of viral replication. There is increasing evidence that HCMV replication is not restricted to early childhood but continues at a lower frequency in (the immunocompetent) adulthood, especially in the eldery [2, 3]. In particular, it is hypothesized that "infectious reactivation in adults is an important driver of transmission of CMV" [2].

The replication cycle of HCMV in HCMV-naive and HCMV-experienced hosts has been analyzed in [15], see also the Review [16]. HCMV replication dynamics were investigated in a population of 30 liver transplant recipients with active HCMV replication. The mean growth rate of virus during infection of HCMV-naive individuals was 1.82 units/day (95% confidence interval (CI), 1.44 − 2.56 units/day), corresponding to a viral doubling time of 0.38 days (95% CI, 0.27 − 0.48 days). Infection of patients with preexisting HCMV immunity was associated with a significantly slower viral growth rate (0.62 units/day; 95% CI, 0.54 − 0.71 units/day) and longer viral doubling time (1.12 days; 95% CI, 0.99 − 1.25 days). In another study the estimated doubling time was significantly longer with a median of 4.3 days and an IQR of 2.5-7.8 days (independent of the immune status) [17].

For a lower bound we estimate the number of days at which the virus is replicating as follows. There is evidence that in early childhood the virus replicates over a long period [18]. Assuming that in early childhood the virus replicates for two years (not necessarily at a stretch, within the first 5 years of infection), that is $365 \cdot 2$ days, we arrive at a lower bound of $365 \cdot 2/4.3 \approx 170$ viral generations during this period. In the elderly host the virus regularly reactivates [19]. In this study 57% of the investigated elderly hosts (which were older than 65 years old) were shedding CMV. On an evolutionary time scale these old hosts might not play a role. Therefore, for a lower bound on the number of viral generations we ignore this time period. Between early childhood and late adulthood the immune system is typically able to keep the virus in check to a large extent. In healthy non-pregnant seropositive women the shedding prevalence was estimated to be 4.7% [3]. Additionally, a shedding prevalence of 3.8% was found in a representative sample of seropositive persons aged 6-49 in the US [20]. Taking the average of both values, we estimate a shedding prevalence of 4.25%. Hence, after infancy and during adulthood, we assume that, on average, the virus is active for $0.0425 \cdot 365 \approx 16$ days each year. This leads to roughly 4 viral generations per year. Assuming again that a typical host is infected for 50 years, we arrive at a lower bound of $170 + 45 \cdot 4 = 350$ viral generations per host life.).

For an upper bound on the number of viral generations within the (on the evolutionary time scale) typical host, we assume that it experiences primary infection in early childhood, followed by a phase of frequent viral replication until age 5 and subsequent 45 years of persistent infection with periodic reactivation. By the above numbers, in the first five years a maximum of roughly $(365/0.38) \cdot 5 \approx 4800$ viral generations should occur. For the subsequent 45 years, we assume the above estimated viral doubling time in the HCMV-experienced host of 1.12 days and that the host carries an active infection roughly half of the time. The latter assumption is motivated by a shedding prevalence of 57% observed in elderly, HCMV-experienced patients, which is supposed to be higher than that of the average adult HCMV-experienced hosts. This yields a total of $45 \cdot 365 \cdot 0.5/1.12 \approx 7730$ viral generations during the second period, and roughly $12530 = 4800 + 7730$ generation in the host's entire life.

#### Primary infection with a single type

On an evolutionary timescale, most hosts are likely to acquire primary HCMV infection via breast milk or during early childhood via saliva or urine, as in regions with limited access to sanitation and hygiene infrastructure and/or high breastfeeding rates, HCMV seroprevalence reaches approximately 85% by the age of one year, see [21] and references therein.

The low virulence of HCMV suggests that at first only a single type can establish the infection in the new host and then further genotypes are transmitted. In general it is reasonable to assume that a host gets infected each time by a different host, as assumed in our model. However, since breast feeding periods are relatively long and it is common that in breast milk several genotypes are detected [22, 23], also transmissions from the same hosts may be relevant. We do not explicitly model this scenario here.

One could account for the transmission of multiple types at primary infection as follows: in addition to the host replacement events defined in the main text and in [24], in which only one type is transmitted from the infecting host to the primary infected host, one could model a mechanism where a random subset of the virus particles in the infecting host are transmitted to the primary infected host. Mathematically, this should yield the same limiting distribution. On the other hand, incorporating extended breastfeeding periods in which the evolution of the viral population within the newborn is coupled with the mother's viral population for an extended time, would likely yield different results.

#### Reinfection

**Aquistion of new genotypes at reinfection** It appears that persistently infected hosts can be reinfected with HCMV despite HCMV-specific humoral and cellular immunity [25].

Women who have been primary infected with CMV recently have been shown to be infected only with a single CMV genotype at the ORFs UL55, UL144, UL146 and UL09 [26]. On the other hand in the majority (15 out of 16) of random samples from CMV-seropositive women (with the same demographic origin) with presumably past primary infections (as the women have been randomly sampled) several genotypes at the ORF UL55 and UL73 have been detected [27]. In [28] genotyping was performed for gB (UL55), gH (UL75) and UL10 for 36 immunocompetent patients during primary infection and in 14 patients experiencing HCMV reactivation/reinfection. In all cases only one gB-gH-UL10 genotype was detected in patients experiencing a primary infection, in contrast in 4 of the 14 non-primary patients a mixed genotype infection was detected. The frequency of mixed infections thus differed significantly between primary and past infection ($p = 0.004$, Fisher's exact test).

These findings indicate that also virions that are transmitted to a host at reinfection may persist in a host and therefore at least part of the diversity observed in hosts may have been introduced at one or several reinfection event.

**Reinfection rates** For simplicity we assume in the model that the rate at which a host is reinfected does not depend on the state of its viral population. Currently, there are no data available on the influence of within-host diversity on the reinfection rate. However, the accumulation of diversity in the virus population within a host may antagonize the host's immune system, boost reactivation and as a consequence increase the reinfection rate.

Two different methods have been used in the literature to assess the frequency of reinfection. In both cases longitudinal samples from patients were taken. In method a) the samples from different time points were sequenced at specific ORFs and the genotypes present in the different samples were compared [29]. The appearance of a new genotype was considered as evidence for reinfection. In method b) the reactivity to antibodies specific for certain genotypes of the ORFs UL55 and UL75 have been analyzed in samples from different time points [30]. The appearance of a new antibody specifity in a host was considered as evidence for reinfection.

Both methods have certain drawbacks. Using deep-sequencing methods it has been shown that genotypes are often present only at very low frequencies and that frequencies of genotypes can vary strongly over time in samples from transplantation patients [31]. At reactivation viral growth is presumably exponential. The virions which initiate the reactivation at the very beginning can grow most and the progeny of these few virions constitute the bulk of the viral population in bodily fluids from which virus is sampled. Therefore the frequencies of the genotypes measured in a single sample might not map well the frequencies of the genotypes in the persistent viral population and often may over-represent a single genotype which is the type of the virion initiating the reactivation. Furthermore, in samples from different time points estimates of genotype frequencies may differ strongly, because the virions sampled may stem from different reactivations. Hence, with method a) it is possible that the genotypes detected at different time points were present in the host over the entire period. This would lead to an overestimation of the reinfection rate. On the other hand, method a) may underestimate the overall reinfection rate since reinfections which do not introduce a new genotype at the sequenced loci go unnoticed.

A problem with using method b) is that a large proportion of hosts (roughly 30%) does not show any genotype-specific antibody reactivity. At least for ORF UL75 the reason cannot be that the hosts react to a different epitope, because in this region the population clusters only into two genotypes. In [30] the authors claim that reactivity to strain specific epitopes was shown to persist on average for 21 months. Hence, it well might be that if a specific genotype is not reactivated the antibody response to that genotype decreases over time, although virions carrying this genotype still persist in the host. A lacking antibody response on the other hand may facilitate the reactivation of the virions with that genotype and renew the antibody response which then is detected in the samples. A reinfection event however need not have occurred to generate this response. This would lead to an overestimation of the reinfection rate with method b).

On the other hand with method b) the reinfection rate might also be underestimated. In [30] an annual reinfection rate has been estimated by first determining the proportions of hosts reinfected in a certain time interval and then dividing this number by the length of the time interval (in time units of years). Using this algorithm the reinfection rate may be underestimated, since hosts who are reinfected with genotypes they had already been infected with earlier may be not recognized as reinfected. This is especially likely if only few genotypes are considered. If for example 20% of the CMV-infected persons are infected only with genotype gH1, 30% of the persons are infected with both genotypes gH1 and gH2 and 50% of the persons are infected only with genotype gH2, a person (so far only infected with genotype gH1) is only in roughly $30 \cdot 0.5$ % + 50% of all reinfections infected with genotype gH2 (if one assumes that a host infected with both genotypes transmits in 50% of the cases genotype gH1 and in 50% of the cases genotype gH2). By determining the proportion of the different genotypes in the population and transmission probabilities of the different genotypes from host infected with several genotypes, one should be able to estimate the reinfection rate quite well also when considering only a few (or a single) ORF (of course provided that one is able to distinguish the occurrence of a new genotype from the reactivation of an already present genotype).

With method a) in 7 of 37 (= 19%) young children attending day care centers evidence for reinfection has been shown within a time frame of sampling between 11 weeks to 38 months with a median of 11 months[29].

In [30] 205 CMV seropositve women and in [32] 149 CMV pregnant women (40 of these with congenitally infected infants and 109 with uninfected newborns) were followed prospectively with method b). In the first study an annual reinfection rate of 10% has been estimated. In the second study an annual reinfection rate of 34% for mothers with congenitally infected infants and an annual rate of 9% for mothers with uninfected newborns have been estimated. Assuming, again, that the from an evolutionary perspective typical host lives for 50 years, we arrive at estimates of $5 \approx 0.09 \cdot 50$ to $17 \approx 0.34 \cdot 50$ reinfections per host life.

It is reasonable that the reinfection rate is smaller than the viral replication rate, because before a host can be reinfected by another host the virus has to be replicated in the reinfecting host first and then this virus has to be transmitted to the reinfected host.

#### Genotypes and mutation between genotypes

For many open reading frames (ORF) a clustering into different genotypes has been described [33]. For some of these ORF it has been shown that different genotypes are differently expressed, for other ORFs the roles of the different genotypes (if they exist) are still open.

An example for an ORF at which genotypes have also different functions is UL 75 encoding the glycoprotein gH. This glycoprotein forms together with the glycoprotein gL (UL115) and either with the proteins encoded by UL128, UL130 and UL131 a pentameric complex or with the glycoprotein gO encoded by UL74 a trimeric complex, both of which are essential for the viral entry into epithelial/endothelial cells and fibroblasts [34, 35, 36]. UL128, UL130 and UL131 are highly conserved regions, but the DNA sequences of UL74, UL75 and UL115 cluster into eight, two and four genotypes, respectively, see [33] and the references therein. Two different antibody responses are triggered by the different versions of glycoprotein gH. The binding site of the neutralizing human antibodies react on two different variants of gH, which are distinguished by at least two amino-acid changes and an amino-acid deletion [37], and called genotype gH1 and gH2.

To arrive at estimates for the mutation rate between genotypes of open reading frames ( not for site-wise mutation rates!) we performed for each locus a phylogenetic analysis to separate the 128 consensus sequences [2] of low-passaged HCMV-samples provided by [9] into two groups. Then we counted the number of amino-acid changes or deletions that are unique for each group. For example for UL75 the genotypes are according to this criterium not only distinguished by the two amino-acid changes and one deletion but by six amino-acid changes and one amino-acid deletion. All six amino-acid changes and the amino-acid deletion are located within a small N-terminal region of a length of 28 amino-acids, see Figure 2. A high differentiation between the two groups of sequences is reflected in a high fixation index $F_{ST}$ of 0.89, calculated according to the formula by [38].

Epithelial/endothelial cells and fibroblasts are cell types in which the virus can replicate effectively [39], and from which virus is released into bodily fluids. As the ORF UL75 is strongly involved in virus entry into these cells, virus sampled from bodily fluids should carry an intact ORF UL75. In samples (from [9]) only the two genotypes gH1 and gH2 are detected, at least on the basis of a consensus sequence. Therefore we will assume in the subsequent analysis that only the two genotypes (which we distinguish by 6 amino-acid changes and one amino-acid deletion) allow a proper virus entry into these cells. As a consequence of our assumption all 6 amino-acid changes and the amino-acid deletion have to occur within a host life.

If we assume in the following that the rate at which amino acids are deleted is the same as the mutation rate per site, the rate at which a change from genotype gH1 to genotype gH2 and vice-verse happens is $\left(\gamma_N \cdot 2 \cdot 10^{-7}\right)^7$, because within a host life on average $\gamma_N$ viral replication events occur, the mutation rate per site was estimated to be $2 \cdot 10^{-7}$ in [40] and all 6 amino-acid changes plus the amino-acid deletion have to occur.

This leads to an upper estimate of $(1.3 \cdot 10^4 \cdot 2 \cdot 10^{-7})^7 \approx 8.0 \cdot 10^{-19}$ for the mutation rate $\mu_N/N$ per virion of the ORF UL75 from genotype gH1 to genotype gH2 and vice versa, see the point “viral replication rate” for an estimate of $\gamma_N$.

Using the same method for the other loci UL4, UL78 and US27 we obtain mutation rate estimates of $5.4 \cdot 10^{-24}$, $1.8 \cdot 10^{-8}$ and $1.2 \cdot 10^{-13}$, respectively. These examples illustrate that it is reasonable to assume that mutation rates between genotypes are very small and in particular mutations happen on a slower time scale than recombination, reinfection and viral reproduction. These estimates are even so small that Assumption **A** 4 might not be fulfilled for the considered loci. However, note, that these rate estimates are based on the assumption that intermediate sequences between genotypes are not viable, which has a strong influence on the estimate of the mutation rate. Further research is necessary to understand how different genotypes arise. In

[2] A consensus sequence is determined as follows: The HCMV DNA from e.g. a urine sample of a HCMV-positive patient is sequenced and the reads are aligned to a reference sequence. Then the consensus sequence is built from this alignment by choosing the most frequent variant at each site.

particular, for highly variable regions the formation of new genotypes could just be a result of genetic drift and mutated genotypes could be as viable as the original ones.

#### Recombination

Several studies have found evidence for frequent recombination in HCMV by analyzing linkage disequilibrium [41, 9]. This also includes recombination between loci subject to our study. Hence it is reasonable to assume that all pairs of loci considered here can be separated by recombination, i.e. that Assumption **R** is fulfilled.

Estimates for recombination rates in HCMV are scarce. The only estimate of a recombination rate for HCMV known to the authors is given in [40]. There, it is estimated that $2N_e r/\text{kb}$ is equal to 2.01, where $N_e$ is the effective population size and $r$ is the per generation recombination rate. In [40] the effective population size is assumed to be 1000. This estimate is based on an estimate of the within host effective population size obtained in [6]. Using this estimate of $N_e$ leads to an estimate of 0.23 crossovers in the whole genome (which is roughly 235 kb long) per viral generation. However, for $N_e$ one should not plug the within host effective population size of the viral population, but the effective population size of the total viral population. This would lead to a greatly reduced recombination rate, at least by a factor $10^{-3}$ which is the inverse of (a lower bound of) the estimated effective population size $M$ of the human (host) population. In the terminology of our model a viral generation takes on average $\frac{1}{\gamma_N}$ time units, so the corrected estimate from [40] would imply $\rho_N = 2.3 \times 10^{-4}\,\gamma_N$. Plugging in the upper bound for $M$, we would get $\rho_N = 2.9 \times 10^{-11}\,\gamma_N$. In both cases this would mean that the recombination rate is much smaller than the viral reproduction rate, which would be in accordance with our model assumptions.

#### Discussion of Assumptions A

We summarized the above discussed parameter estimates in Table 1.

**Assumption A 2** By the estimates, it seems reasonable to believe that the mutation rate and the recombination rate are of a lower order than the viral replication rate, i.e.

$$\mu_N, \rho_N \ll \gamma_N.$$

Whether or not the viral replication rate is much greater than the effective population size of the persistent population size, i.e. whether or not

$$N \ll \gamma_N$$

holds, is uncertain: On the one hand, the physical size of the persistent population could be of the same order or even higher than $\gamma_N$ by the estimates above. On the other hand, effective population sizes are often much smaller than physical ones.

**Assumption A 3** The estimates for the rate of mutation between genotypes $\mu_N$ lie between $(\gamma_N \cdot 2 \cdot 10^{-7})^3 \cdot N$ and $(\gamma_N \cdot 2 \cdot 10^{-7})^9 \cdot N$. Only for UL78 (where only three amino acid changes uniquely separate genotypes) there is a parameter combination within the estimated parameter ranges such that $\rho_N$ is not much greater than $\mu_N$.

**Assumption A 4** This assumption is not fulfilled, if the mutation rate is estimated as described in the subsection "Genotypes and mutation between genotypes", which relies on the assumption that intermediate sequences between genotypes are not viable and hence all mutations that distinguish two genotypes have to arise within one host life. Note, however, that our data analysis does not rely on Assumption A 4, because by considering bi-allelic loci we condition on the event that the genotypes are present in the viral population at non-trivial frequencies.

**Assumption A 1** This assumption states that the relationship between the number of viral generations per host life and the number of reinfection events per host life is asymptotically constant and given by a parameter $\theta$. In the main text, we estimated this parameter by $\hat{\theta} = 0.147$ which would mean that, on average, roughly every 7 viral generations a reinfection event occurs. In other words, if the evolutionarily typical host lives for 50 years, then, in the above described parameter ranges, it would experience between 51 and $1.9 \times 10^3$ reinfections. This is much higher than previous estimates in the literature which range from 5 to 17 reinfections per host life.

These previous estimates are, however, subject to great uncertainty as explained above. They typically only count the *successful* acquisition of *new* genotypes at a certain small set of considered loci as a reinfection. In our model also those reinfection events at which the reinfected host acquires no new genotypes are counted, as well as those at which new genotypes are acquired but quickly lost due to random genetic drift.

Table 1: Overview of parameter regimes inferred from the literature. The estimates of the rate $\lambda_N$ are based on estimates available in the literature and the estimates of the rate $\hat{\theta}\gamma_N$ are the estimates for the reinfection rate based on estimates of $\gamma_N$ and the inferred value $\hat{\theta}$.

| Symbol | Interpretation | Lower estimate | Upper estimate | Unit |
|---|---|---|---|---|
| $N$ | persistent pop. size | | $7 \times 10^3 - 2 \times 10^5$ | virions |
| $M$ | host pop. size | $10^3 - 10^4$ | $8 \times 10^9$ | hosts |
| $\gamma_N$ | viral replication rate | 350 | $1.3 \times 10^4$ | vir. generations/host life |
| $\lambda_N$ | reinfection rate | 5 | 17 | reinfections/host life |
| $\hat{\theta}\gamma_N$ | implied reinfection rate | 51 | $1.9 \times 10^3$ | reinfections/host life |
| $\mu_N/N$ | mutation rate | $5.4 \cdot 10^{-24}$ | $1.8 \cdot 10^{-8}$ | mutations/(virion per host life) |
| $\rho_N$ | recombination rate | $2.9 \times 10^{-11}\gamma_N$ | $2.3 \times 10^{-4}\gamma_N$ | recombinations/host life |

### 2 Age groups and sample determination

Much of the literature on HCMV focuses on specific host populations, e.g. newborns, pregnant women, or transplant recipients. The goal of this paper is to model the total population of persistently infected hosts in which the virus is actively replicated rather than a specific subgroup. Since hosts get reinfected over time, older hosts might carry more diverse viral populations than younger ones (and this pattern can also be observed in the literature, see e.g. [42]). For this reason we aimed for a sample that is representative (with respect to age) of this total population.

In order to account for the age structure and mimic a random sample from the population of persistently HCMV infected hosts, we use stratified sampling. We consider the age groups "infants" (0-4.9 years), "children and adults" ("c &a" for short, 5-59.9 years) and "seniors" (60+ years). For simplicity (and data availability) we assume that HCMV is being replicated in a host mainly if the host is shedding HCMV DNA detectable by PCR. For our study we want to mimic a sample from the population of HCMV infected hosts in which the virus is actively being replicated. To generate the sample, we need to estimate the percentages of the age groups "infants", "c & a" and "seniors" within the HCMV shedding population in Austria. That is, we want to estimate

$$\mathbb{P}\big(\text{ infant } \big| \text{ shedding }\big), \mathbb{P}\big(\text{ c\&a } \big| \text{ shedding }\big) \text{ and } \mathbb{P}\big(\text{ senior } \big| \text{ shedding }\big). \tag{1}$$

To our knowledge, there are no estimates for these quantities in the literature. There are, however, estimates for sero- and shedding prevalences which we can use in combination with demographic information from census data to obtain estimates for (1).

**Infants.** Bayes' Theorem implies that

$$\mathbb{P}\big(\text{ infant } \big| \text{ shedding }\big) = \frac{\mathbb{P}\big(\text{ infant and shedding }\big)}{\mathbb{P}\big(\text{ shedding }\big)} = \frac{\mathbb{P}\big(\text{ shedding } \big| \text{ infant }\big)\mathbb{P}\big(\text{ infant }\big)}{\mathbb{P}\big(\text{ shedding }\big)}. \tag{2}$$

From the 2011 Austrian census data [43], which were the most recent officially published census data at the time the study was designed, we estimate $\mathbb{P}\big(\text{ infant }\big)$ to be roughly 0.047. Estimates for $\mathbb{P}(\text{ shedding } | \text{ infant })$ for the Austrian population are not available to our knowledge, hence we use shedding rates observed in US day care centers which are given in [18], Table A1, Category II. Note, however, that the shedding rates in [18] were determined using culture methods as opposed to PCR and hence may underestimate the shedding rate as we defined it above. On the other hand, the shedding rates in [18], may overestimate the shedding rate within the general population of 0-5 year olds since they were observed in day care centers where the exposure to HCMV may be higher than in the general population. Thus we take the average in [18], Table A1, Category II to arrive at an estimate of 0.31 for $\mathbb{P}\big(\text{ shedding } \big| \text{ infant }\big)$. The only missing value, $\mathbb{P}\big(\text{ shedding }\big)$, is computed further below.

**C & A.** We calculate

$$\mathbb{P}\big(\text{ c \& a }\big|\text{ shedding }\big)$$
$$= \frac{\mathbb{P}\big(\text{ c \& a }\big)\mathbb{P}\big(\text{ seropositive }\big|\text{ c \& a }\big)\mathbb{P}\big(\text{ shedding }\big|\text{ seropositive and c \& a }\big)}{\mathbb{P}\big(\text{ shedding }\big)}. \tag{3}$$

Again, the 2011 Austrian census yields $\mathbb{P}\big(\text{ c \& a }\big) \approx 0.72$. According to seroprevalence estimates for the German adult population given in [44], a value of 0.5 could be a reasonable estimate for $\mathbb{P}\big(\text{ seropositive }\big|\text{ c \& a }\big)$. There are two estimates for $\mathbb{P}\big(\text{ shedding }\big|\text{ seropositive and c \& a}\big)$ in the literature. In one study [3], the (salivary and urinary) shedding prevalence observed in 102 immunocompetent seropositive nonpregnant female college students, aged 18 to 30, was 4.7%. Another study [20] found a urinary shedding prevalence of 3.8% in seropositive persons aged 6 to 49. Taking the average of both values we estimate $\mathbb{P}\big(\text{ shedding }\big|\text{ seropositive and c \& a}\big) \approx 4.25\%$.

**Seniors.** An analgous calculation as for the other age groups yield

$$\mathbb{P}\big(\text{ senior }\big|\text{ shedding }\big)$$
$$= \frac{\mathbb{P}\big(\text{ senior }\big)\mathbb{P}\big(\text{ seropositive }\big|\text{ senior }\big)\mathbb{P}\big(\text{ shedding }\big|\text{ seropositive and senior }\big)}{\mathbb{P}\big(\text{ shedding }\big)}. \tag{4}$$

By the 2011 Austrian census, $\mathbb{P}\big(\text{ senior }\big)$ lies around 0.23. The numbers in [44] suggest a seroprevalence of around 70% in this age group, i.e. $\mathbb{P}\big(\text{ seropositive }\big|\text{ senior }\big) = 0.7$. In [19], shedding of HCMV in 11 elderly seropositive individuals was studied. The shedding prevalence was 0.57 which we choose as an estimate for $\mathbb{P}\big(\text{ shedding }\big|\text{ seropositive and senior }\big)$.

Finally, $\mathbb{P}(\text{ shedding })$ can be calculated by computing the sum of the numerators in (2), (3) and (4). Taking everything together, we get

$$\mathbb{P}\big(\text{ infant }\big|\text{ shedding }\big) \approx 0.12$$
$$\mathbb{P}\big(\text{ c \& a }\big|\text{ shedding }\big) \approx 0.12$$
$$\mathbb{P}\big(\text{ senior }\big|\text{ shedding }\big) \approx 0.76.$$

We used these proportions as a rough orientation for the age structure of the sample. Table 2 gives the age proportions in our sample.

Table 2: Age distribution in the sample.

| | infant | children & adults | seniors |
|---|---|---|---|
| Age | 0-4.99 | 5-59.99 | 60- |
| Number of samples | 11 | 9 | 44 |
| Relative frequency | 0.17 | 0.14 | 0.69 |

Table 3: Oligos for amplicon deep sequencing

| **Target** | **Size [bp]** | **Annealing T [°C]** | **Primer ID** | **5' - 3' sequence** |
|---|---|---|---|---|
| UL4 | 420-426 | 50 | UL4_fwC | TGCGCTAACTACTCAGGGACC |
| | | | UL4_fwT | TGCGCTAATTACTCAGGGACC |
| | | | UL4_rv | TACACAGTCAGAGTAGCCTAG |
| UL75 | 341-344 | 49 | UL75_fw | CATCGAGGCATATGGAATACA |
| | | | UL75_rv | GTGTAACGCCAACCACCAC |
| UL78 | 300 | 50 | UL78_fw | TGGTGGGTATGTCCCCTTCT |
| | | | UL78_rv | ATACCCTTGGACAACATGGT |
| US27 | 302 | 51 | UL78_fw | CTGATGGCGTACACGTACAA |
| | | | UL78_rv | GTCCACATGTCCTTCCTCAT |

Table 4: Genotype frequencies $x_\ell^i$

| | **frequency of GT 1 at locus** | | | |
|---|---|---|---|---|
| **individual no.** | **UL4** | **UL75** | **UL78** | **US27** |
| 1 | 0.167152221412964 | 1 | 0 | 1 |
| 2 | 0 | 0 | 0 | 1 |
| 3 | 1 | 0 | 1 | 0 |
| 4 | 0 | 0 | 0 | 0 |
| 5 | 0 | 1 | 1 | 1 |
| 6 | 1 | 1 | cov too low | 1 |
| 7 | 1 | 0 | 1 | 1 |
| 8 | 1 | 0 | 0 | 1 |
| 9 | 1 | 1 | 1 | 1 |
| 10 | 1 | 0 | 1 | 1 |
| 11 | 0 | 0 | 1 | 1 |
| 12 | 0 | 0 | 1 | 0 |
| 13 | 1 | 1 | nd | 1 |
| 14 | 0.757968932387537 | 0.499775684163302 | 0.42040857605178 | 1 |
| 15 | 0 | no cov | 1 | 1 |
| 16 | 0 | 0 | 1 | 0 |
| 17 | 1 | 0 | 1 | 0 |
| 18 | nd | 1 | 1 | 1 |
| 19 | 1 | nd | 1 | 1 |
| 20 | 1 | 0 | nd | nd |
| 21 | nd | 1 | 0 | 0 |
| 22 | 0 | 0 | 1 | 1 |
| 23 | nd | nd | 1 | nd |
| 24 | 1 | 1 | 0 | 1 |
| 25 | 1 | 1 | 0 | 1 |
| 26 | 0.341891008498247 | 1 | 0 | 0 |
| 27 | 0.07223275428483 | 0.114893617021277 | 1 | 0 |
| 28 | 1 | 0.474247270320944 | 0 | 0.701960784313725 |
| 29 | nd | 0 | 1 | nd |
| 30 | 0.419254954268293 | 0 | 0.867488789237668 | 0.609177215189873 |
| 31 | nd | 0 | 0 | nd |
| 32 | 0 | 0.130300096805421 | 0.126578947368421 | 0 |
| 33 | 1 | 0 | cov too low | nd |
| 34 | 1 | 0 | 1 | 1 |
| 35 | 1 | 0 | 0 | 0 |
| 36 | 1 | 1 | 0 | 0 |
| 37 | nd | 1 | 1 | 0 |
| 38 | 1 | nd | 0 | nd |
| 39 | 0 | 0 | 1 | 1 |
| 40 | 1 | 0 | 0 | 1 |
| 41 | nd | nd | cov too low | 1 |
| 42 | 1 | 1 | 1 | 0 |
| 43 | 1 | 0.47674214498859 | 1 | 0 |
| 44 | cov too low | nd | nd | 0.695571955719557 |
| 45 | 0 | 1 | 0 | 1 |
| 46 | 0 | 1 | 1 | 0 |
| 47 | 1 | 1 | 0 | 1 |
| 48 | 1 | 1 | 0.183295688893253 | 1 |
| 49 | 1 | 0 | 1 | 1 |
| 50 | 1 | 0 | 1 | 0.0247749227837287 |
| 51 | 1 | 1 | 0 | 1 |
| 52 | 1 | 1 | 1 | 1 |
| 53 | 1 | 0.610683666739943 | 1 | 1 |
| 54 | 1 | 0.53448275862069 | 1 | 0.558224696059763 |
| 55 | 1 | 1 | 0 | 1 |
| 56 | 0 | 1 | 1 | 1 |
| 57 | 0 | 0 | 0 | 0 |
| 58 | 0 | 0 | 1 | 1 |
| 59 | 0 | 0.613938053097345 | 0.344262295081967 | 1 |
| 60 | nd | cov too low | 1 | 0.719928825622776 |
| 61 | 1 | 1 | 0 | 1 |
| 62 | 1 | 0 | 0 | 1 |
| 63 | 0.197141727390838 | 0.220261171189553 | 0.85122679475529 | 0.916632274040446 |
| 64 | 0.508576726935559 | 0.0230872483221476 | 1 | 0.761140819964349 |

Table 5: Infection Patterns. *1*: only genotype 1 present, *2*: only genotype 2 present, *m* = *"mixed"*: both genotypes present, *nd:* no PCR product, *cov too low:* average coverage < 100 reads.

| | infection pattern at locus | | | |
|---|---|---|---|---|
| **individual no.** | **UL4** | **UL75** | **UL78** | **US27** |
| 1 | m | 1 | 2 | 1 |
| 2 | 2 | 2 | 2 | 1 |
| 3 | 1 | 2 | 1 | 2 |
| 4 | 2 | 2 | 2 | 2 |
| 5 | 2 | 1 | 1 | 1 |
| 6 | 1 | 1 | cov too low | 1 |
| 7 | 1 | 2 | 1 | 1 |
| 8 | 1 | 2 | 2 | 1 |
| 9 | 1 | 1 | 1 | 1 |
| 10 | 1 | 2 | 1 | 1 |
| 11 | 2 | 2 | 1 | 1 |
| 12 | 2 | 2 | 1 | 2 |
| 13 | 1 | 1 | nd | 1 |
| 14 | m | m | m | 1 |
| 15 | 2 | no cov | 1 | 1 |
| 16 | 2 | 2 | 1 | 2 |
| 17 | 1 | 2 | 1 | 2 |
| 18 | nd | 1 | 1 | 1 |
| 19 | 1 | nd | 1 | 1 |
| 20 | 1 | 2 | nd | nd |
| 21 | nd | 1 | 2 | 2 |
| 22 | 2 | 2 | 1 | 1 |
| 23 | nd | nd | 1 | nd |
| 24 | 1 | 1 | 2 | 1 |
| 25 | 1 | 1 | 2 | 1 |
| 26 | m | 1 | 2 | 2 |
| 27 | m | m | 1 | 2 |
| 28 | 1 | m | 2 | m |
| 29 | nd | 2 | 1 | nd |
| 30 | m | 2 | m | m |
| 31 | nd | 2 | 2 | nd |
| 32 | 2 | m | m | 2 |
| 33 | 1 | 2 | cov too low | nd |
| 34 | 1 | 2 | 1 | 1 |
| 35 | 1 | 2 | 2 | 2 |
| 36 | 1 | 1 | 2 | 2 |
| 37 | nd | 1 | 1 | 2 |
| 38 | 1 | nd | 2 | nd |
| 39 | 2 | 2 | 1 | 1 |
| 40 | 1 | 2 | 2 | 1 |
| 41 | nd | nd | cov too low | 1 |
| 42 | 1 | 1 | 1 | 2 |
| 43 | 1 | m | 1 | 2 |
| 44 | cov too low | nd | nd | m |
| 45 | 2 | 1 | 2 | 1 |
| 46 | 2 | 1 | 1 | 2 |
| 47 | 1 | 1 | 2 | 1 |
| 48 | 1 | 1 | m | 1 |
| 49 | 1 | 2 | 1 | 1 |
| 50 | 1 | 2 | 1 | m |
| 51 | 1 | 1 | 2 | 1 |
| 52 | 1 | 1 | 1 | 1 |
| 53 | 1 | m | 1 | 1 |
| 54 | 1 | m | 1 | m |
| 55 | 1 | 1 | 2 | 1 |
| 56 | 2 | 1 | 1 | 1 |
| 57 | 2 | 2 | 2 | 2 |
| 58 | 2 | 2 | 1 | 1 |
| 59 | 2 | m | m | 1 |
| 60 | nd | cov too low | 1 | m |
| 61 | 1 | 1 | 2 | 1 |
| 62 | 1 | 2 | 2 | 1 |
| 63 | m | m | m | m |
| 64 | m | m | 1 | m |

Table 6: Empirical frequencies of the infection patterns. These are the value counts of table 5, excluding the 16 rows containing missing values. *1*: only genotype 1 present, *2*: only genotype 2 present, $m =$ *"mixed"*: both genotypes present

| **infection pattern at locus** | | | | **Empirical Frequency** |
|---|---|---|---|---|
| **UL4** | **UL75** | **UL78** | **US27** | |
| 1 | 1 | 2 | 1 | 6 |
| 1 | 2 | 1 | 1 | 4 |
| 2 | 2 | 1 | 1 | 4 |
| 1 | 2 | 1 | 2 | 3 |
| 1 | 2 | 2 | 1 | 3 |
| 2 | 2 | 1 | 2 | 2 |
| 2 | 2 | 2 | 2 | 2 |
| 2 | 1 | 1 | 1 | 2 |
| 1 | 1 | 1 | 1 | 2 |
| m | m | m | 1 | 2 |
| 1 | m | 1 | 1 | 1 |
| 2 | 1 | 2 | 1 | 1 |
| 2 | 1 | 1 | 2 | 1 |
| 1 | 1 | m | 1 | 1 |
| m | 1 | 2 | 1 | 1 |
| 1 | m | 1 | m | 1 |
| 2 | m | m | 1 | 1 |
| 1 | 1 | 1 | 2 | 1 |
| 1 | m | 1 | 2 | 1 |
| 1 | m | 2 | m | 1 |
| 1 | 1 | 2 | 2 | 1 |
| 1 | 2 | 2 | 2 | 1 |
| 2 | m | m | 2 | 1 |
| m | 2 | m | m | 1 |
| 2 | 2 | 2 | 1 | 1 |
| 2 | m | 1 | 2 | 1 |
| m | 1 | 2 | 2 | 1 |
| m | 2 | 1 | m | 1 |
| | | | | $\sum = 48$ |

Figure 1: Signature positions for UL4. Arrows in the subfamily logo plot at the top indicate the positions used for genotyping. Sequence IDs in the left column correspond to patient IDs. Multiple occurrences of the same patient ID indicate that multiple sequences were found in the material sampled from that patient.

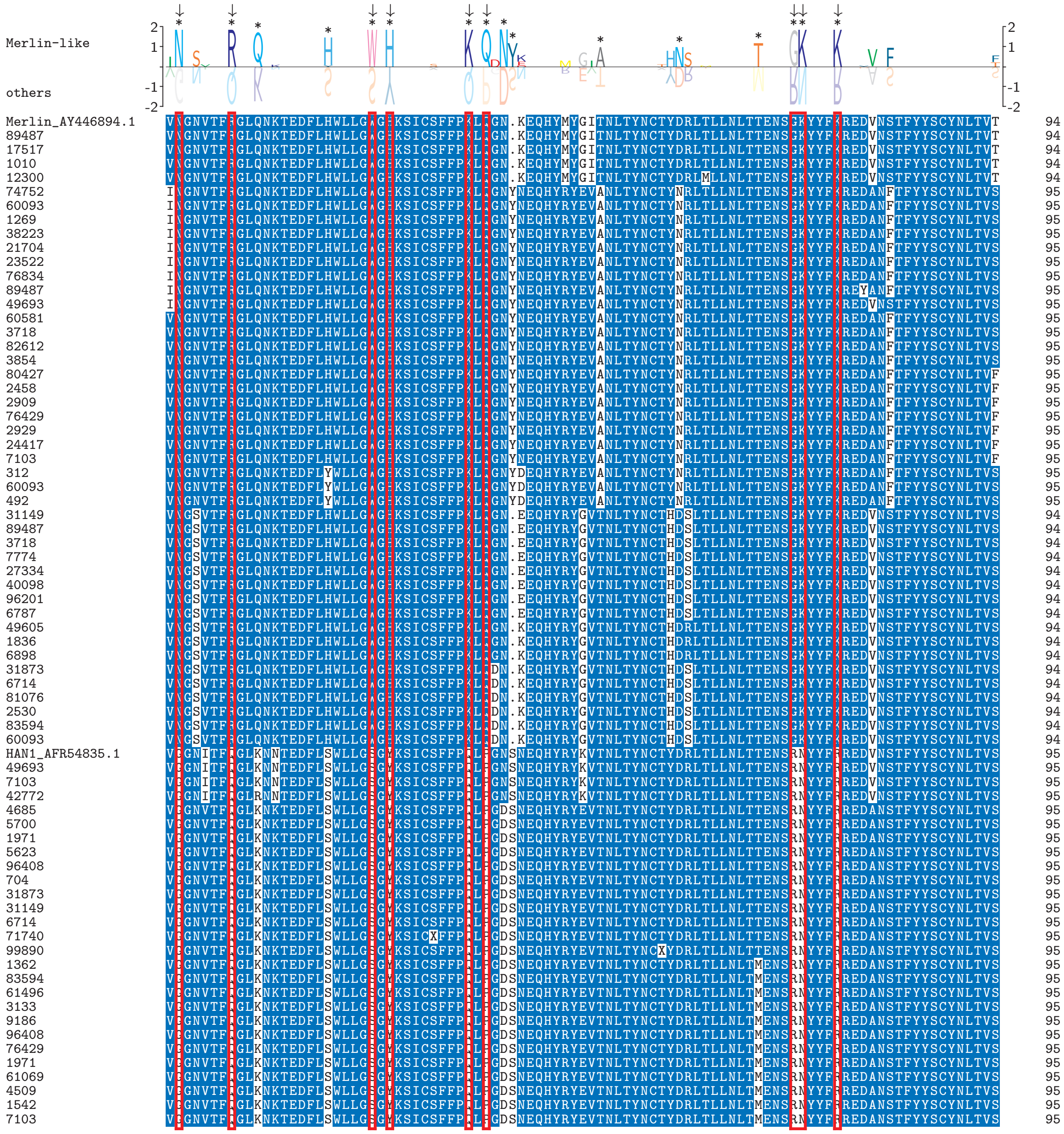

Figure 2: Signature positions for UL75. Arrows in the subfamily logo plot at the top indicate the positions used for genotyping. Sequence IDs in the left column correspond to patient IDs. Multiple occurrences of the same patient ID indicate that multiple sequences were found in the material sampled from that patient.

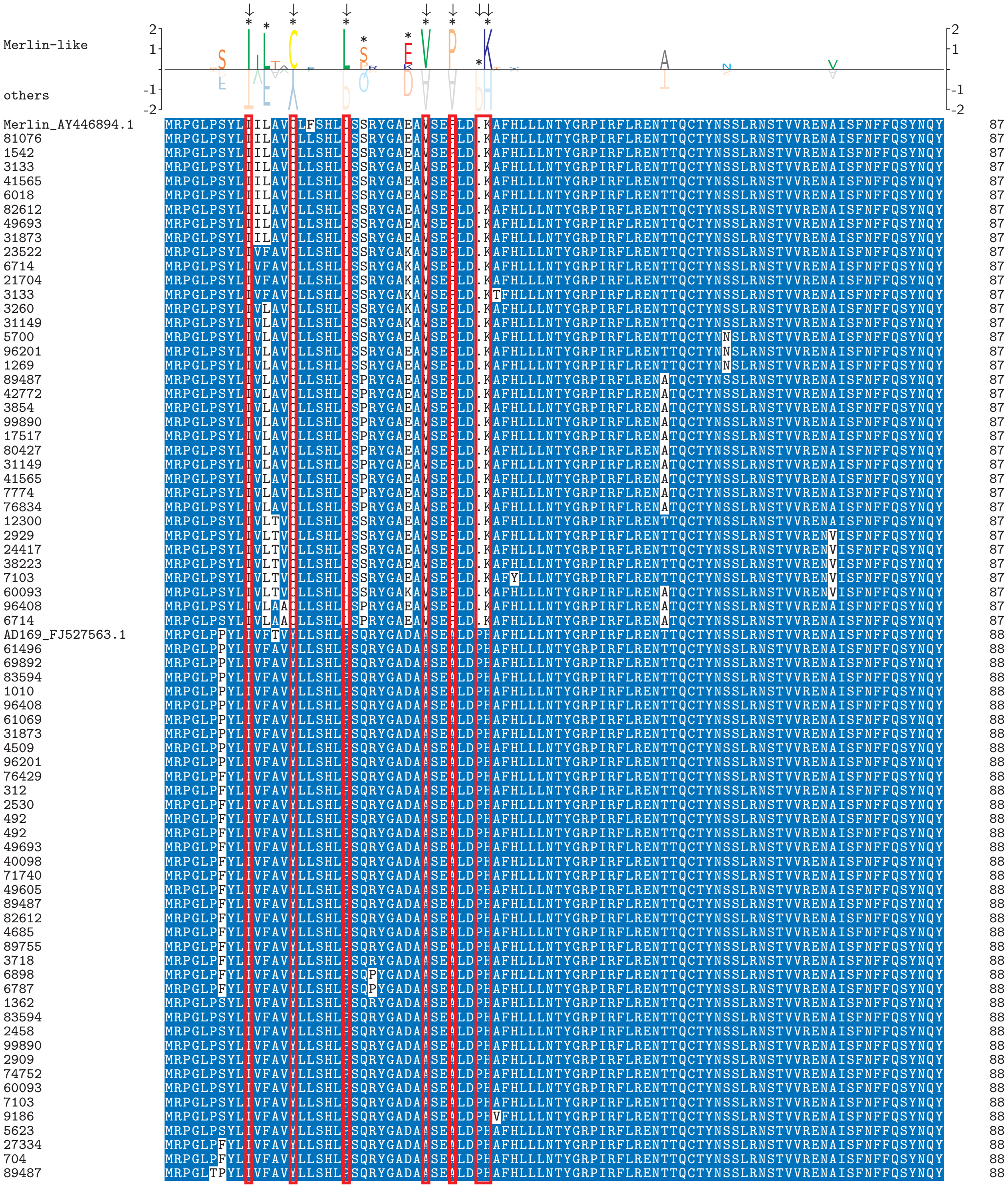

Figure 3: Signature positions for UL78. Arrows in the subfamily logo plot at the top indicate the positions used for genotyping. Sequence IDs in the left column correspond to patient IDs. Multiple occurrences of the same patient ID indicate that multiple sequences were found in the material sampled from that patient. One sequence from patient 96408 did not cluster as clearly into the “Merlin-like” or “other” genotype. It was counted as “Merlin-like” due to its slighty higher similarity with that group.

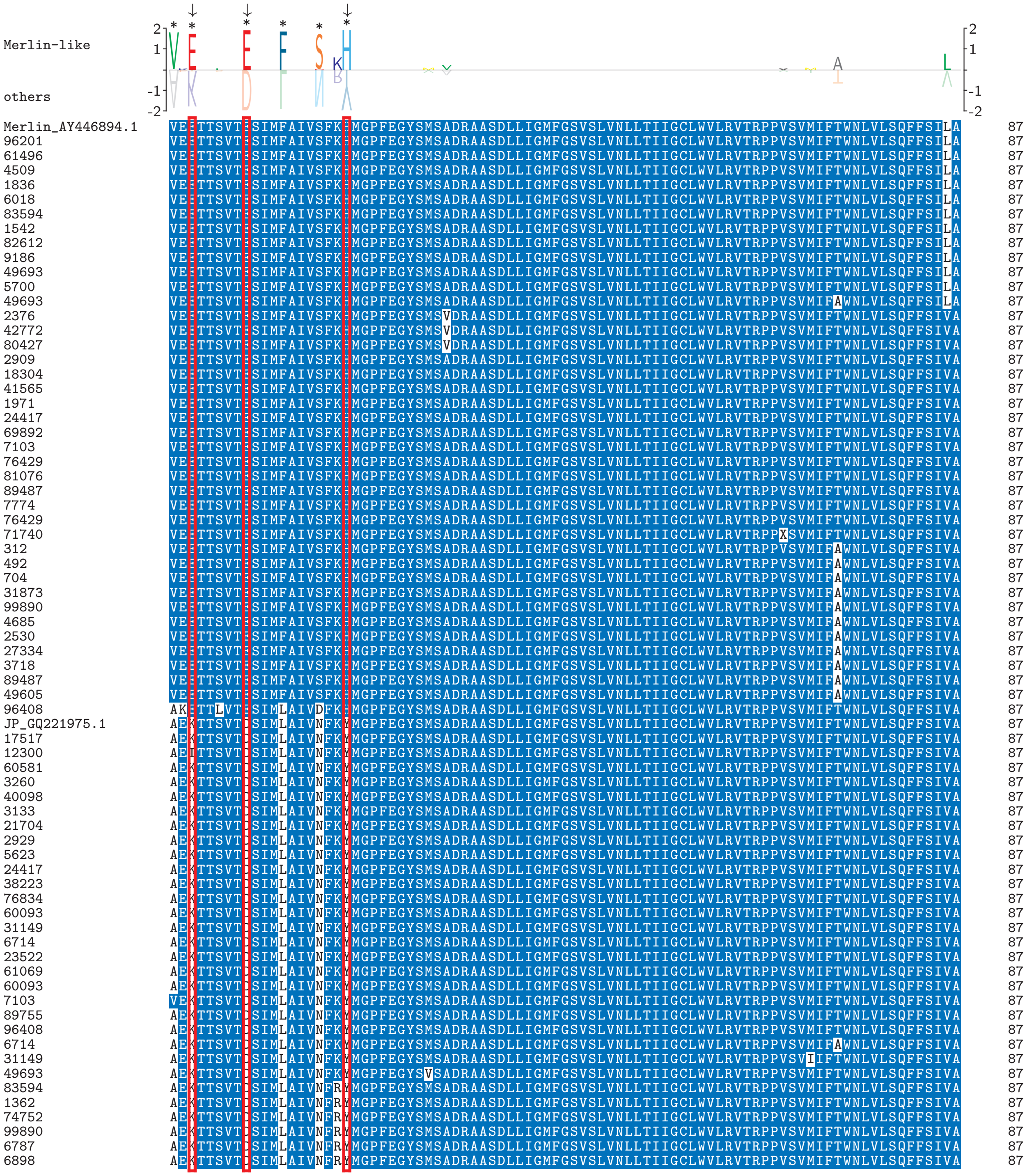

Figure 4: Signature positions for US27. Arrows in the subfamily logo plot at the top indicate the positions used for genotyping. Sequence IDs in the left column correspond to patient IDs. Multiple occurrences of the same patient ID indicate that multiple sequences were found in the material sampled from that patient.

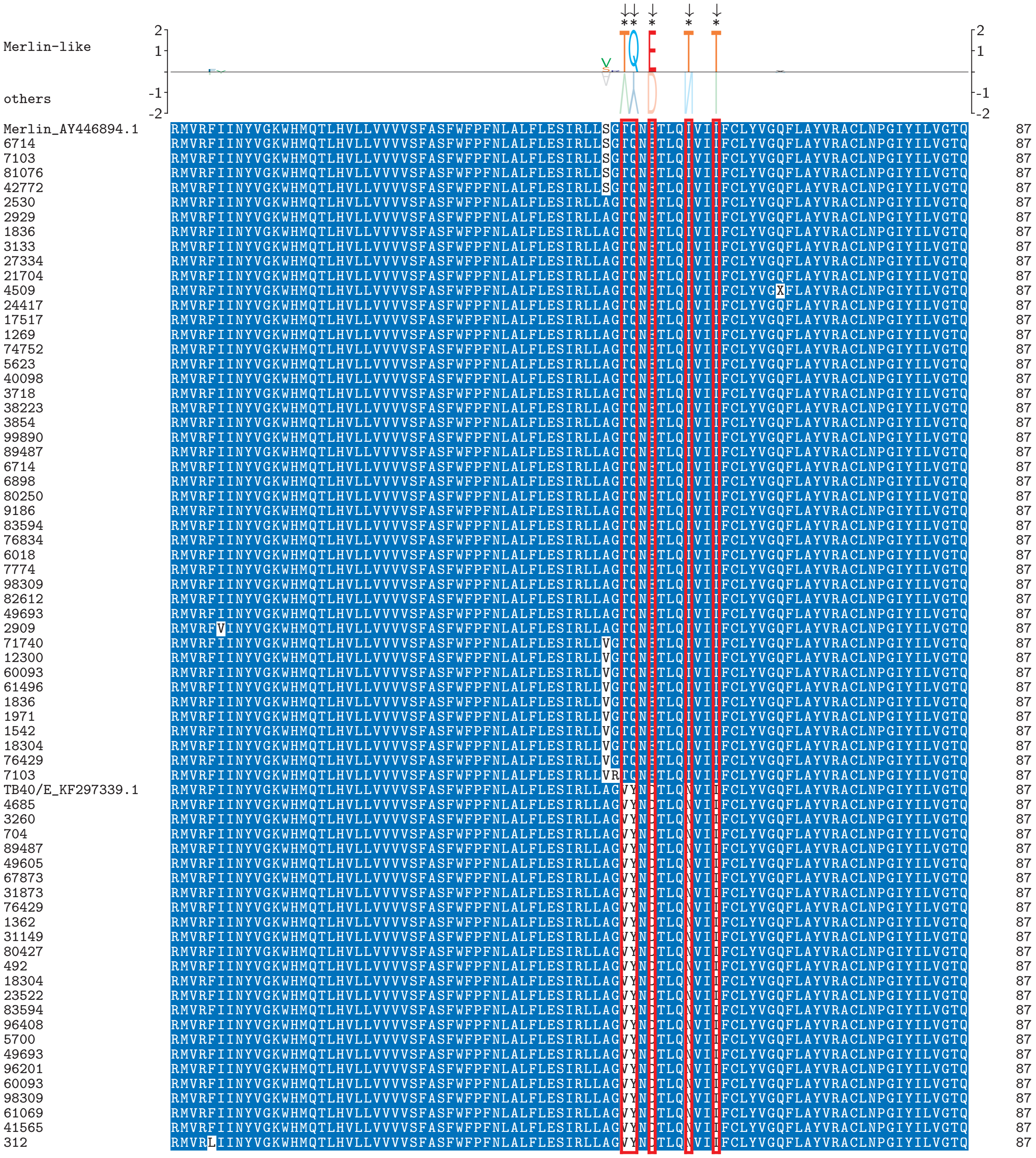

Figure 5: Phylogenetic tree inferred for sequences from [9] at ORF UL4.

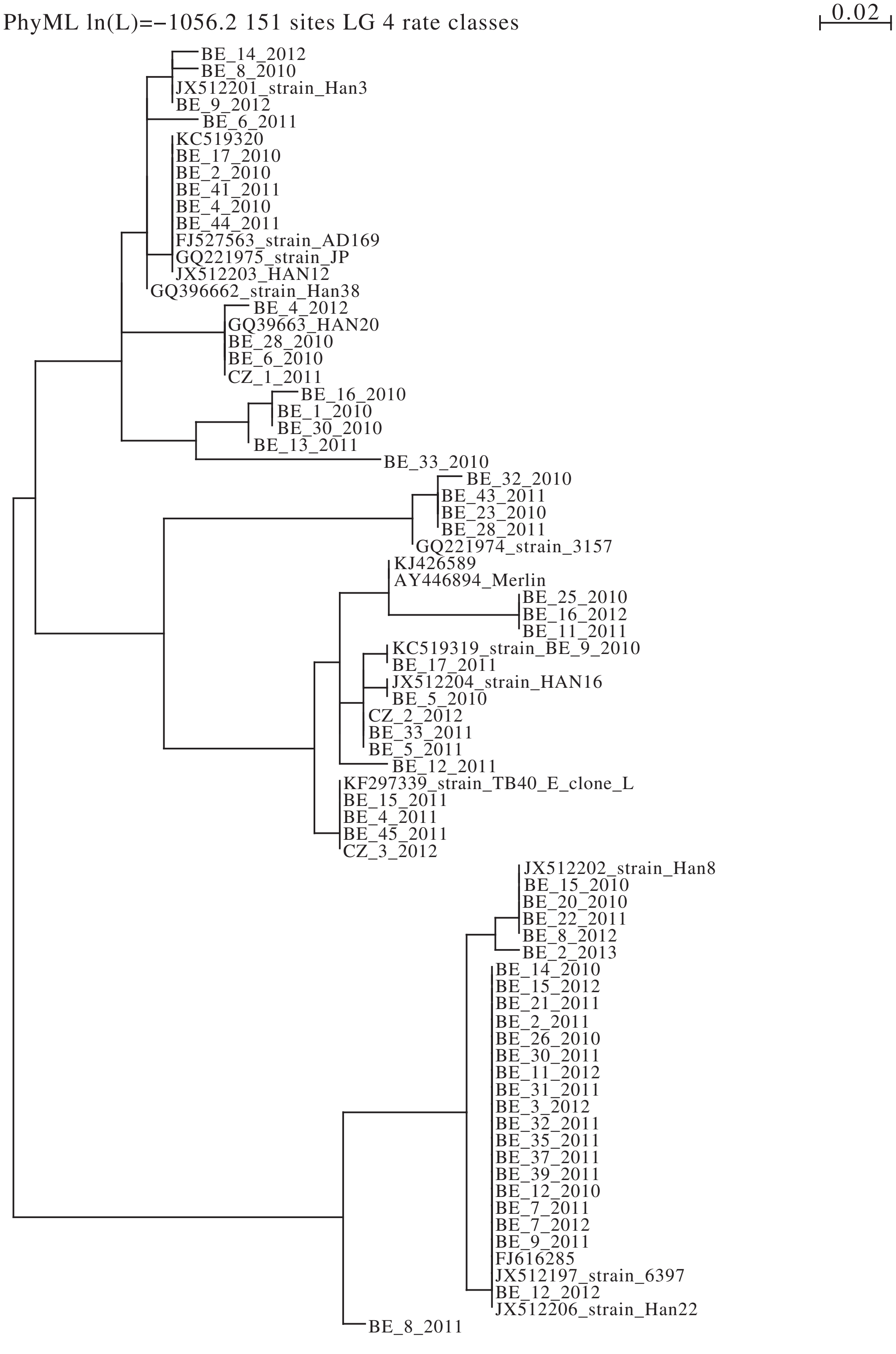

Figure 6: Phylogenetic tree inferred for sequences from [9] at ORF UL75.

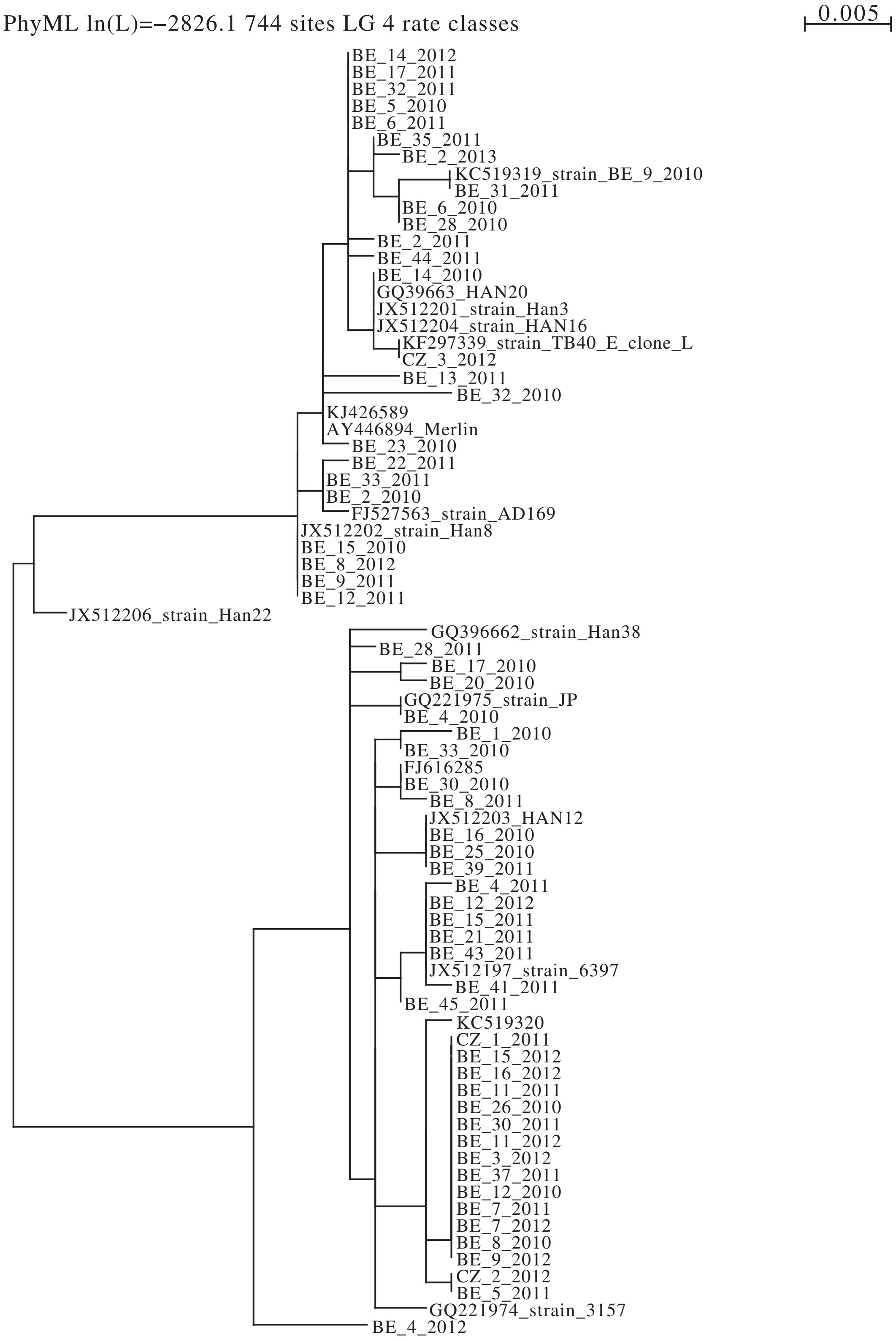

Figure 7: Phylogenetic tree inferred for sequences from [9] at ORF UL78.

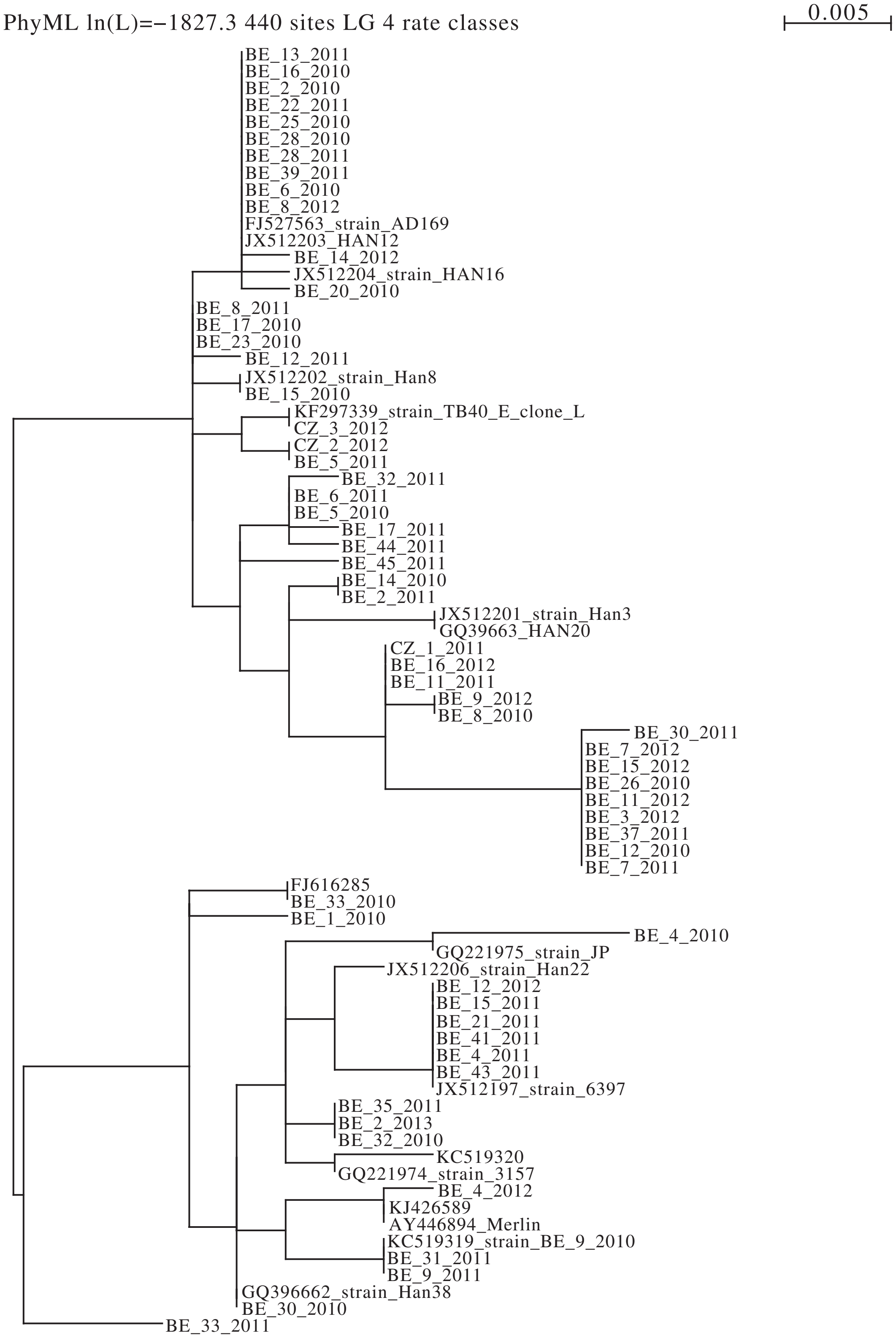

Figure 8: Phylogenetic tree inferred for sequences from [9] at ORF US27.

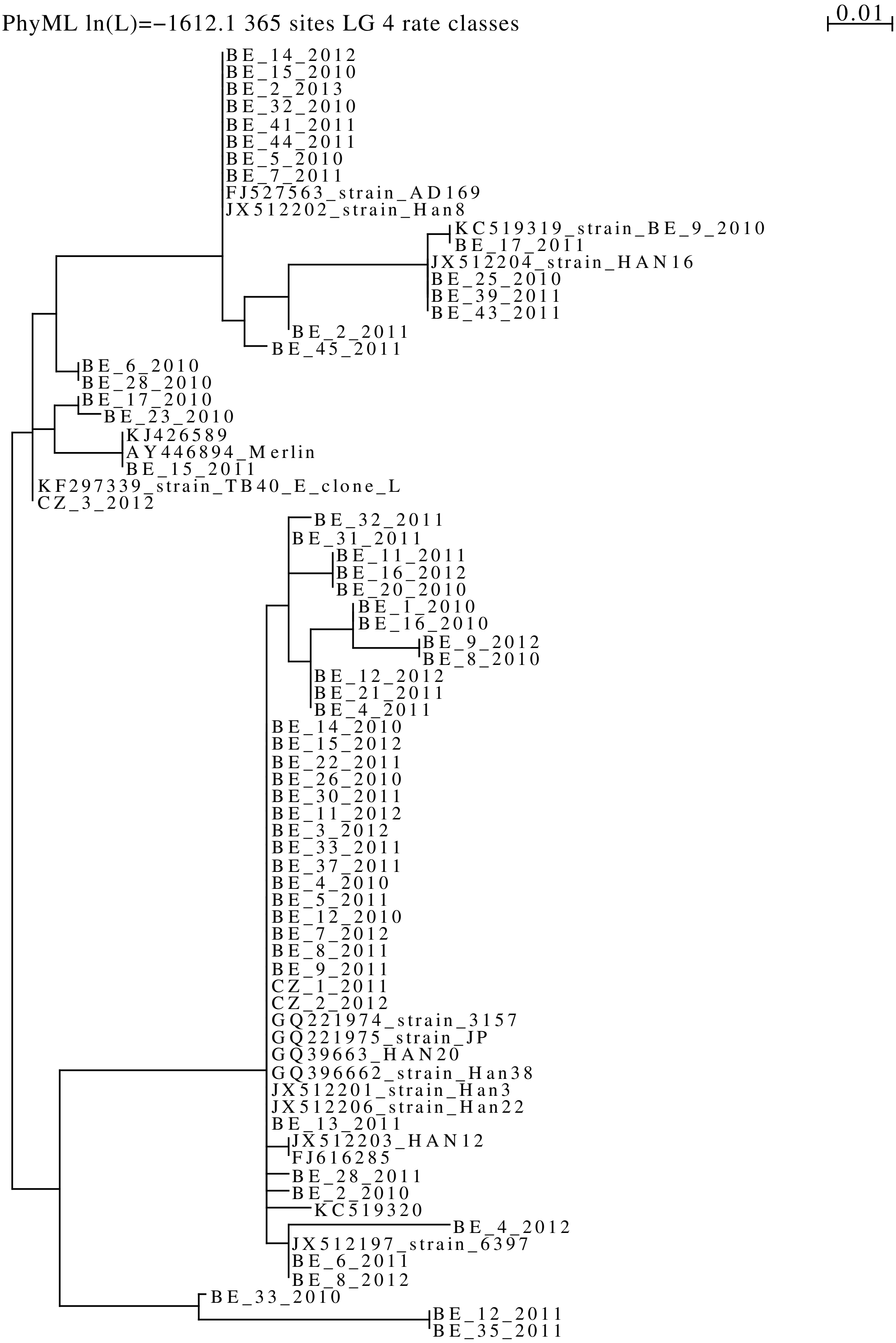

Figure 9: Phylogenetic tree at ORF UL4 inferred from the Austrian sample subject to to our article.

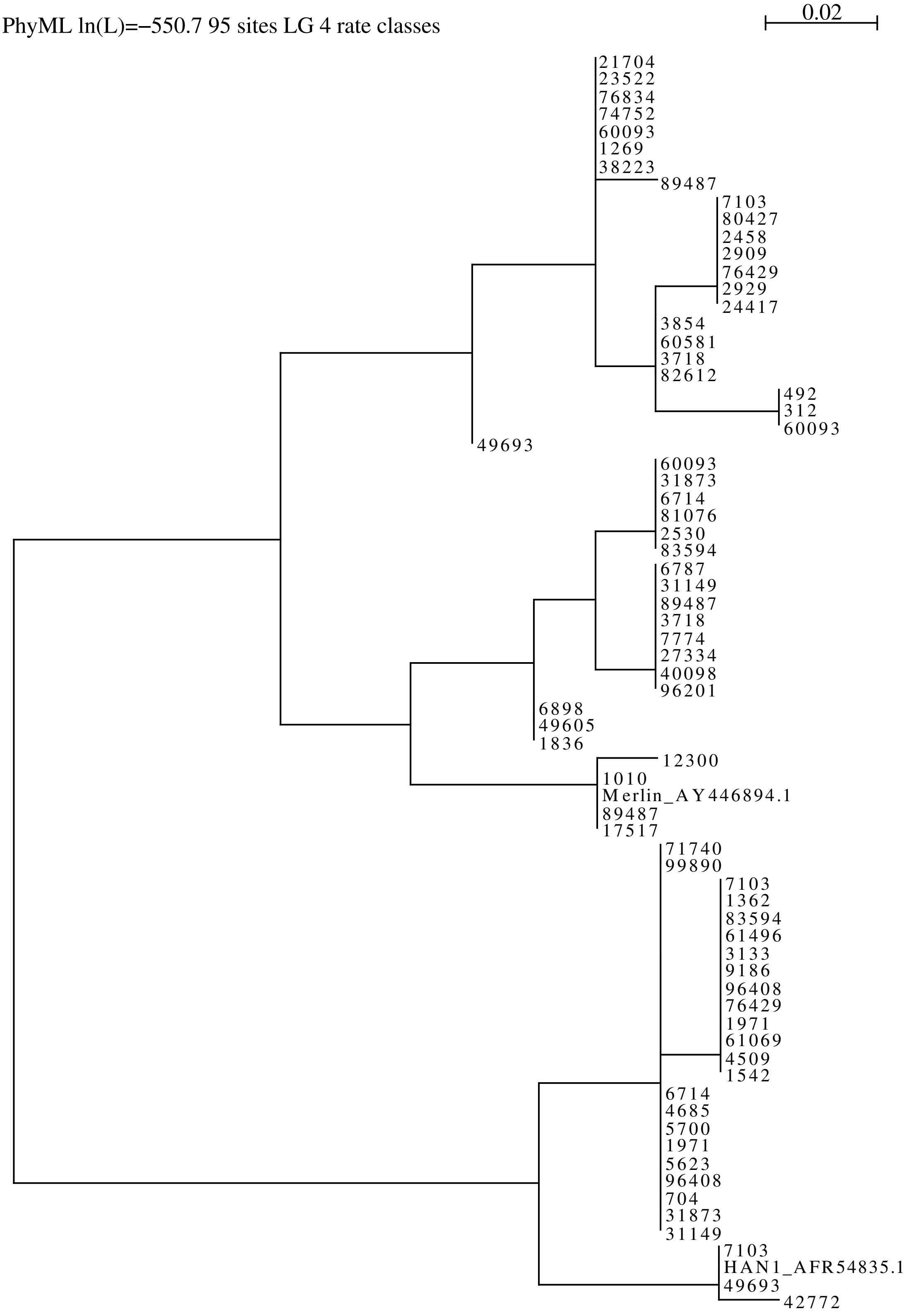

Figure 10: Phylogenetic tree at ORF UL75 inferred from the Austrian sample subject to to our article.

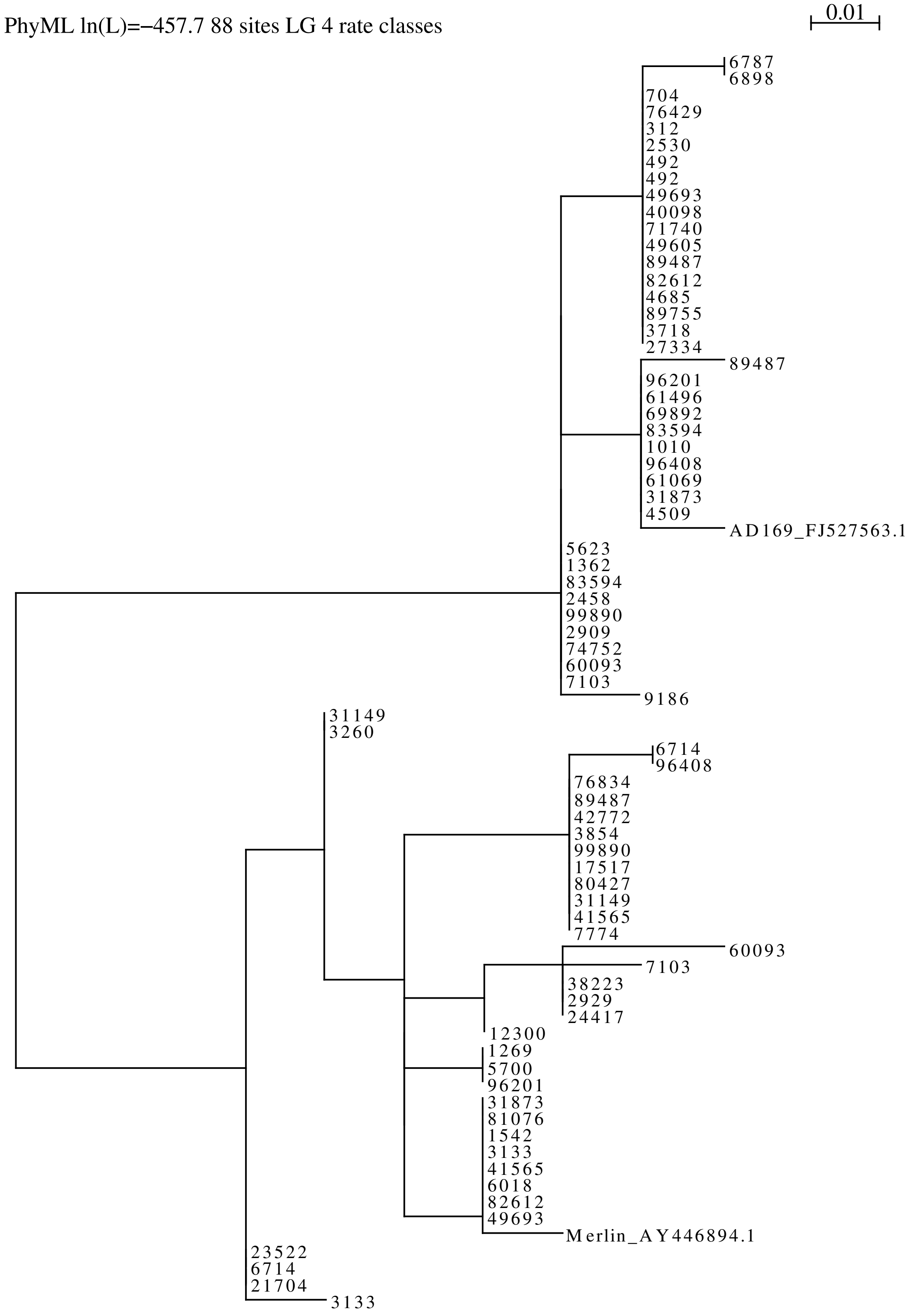

Figure 11: Phylogenetic tree at ORF UL78 inferred from the Austrian sample subject to to our article.

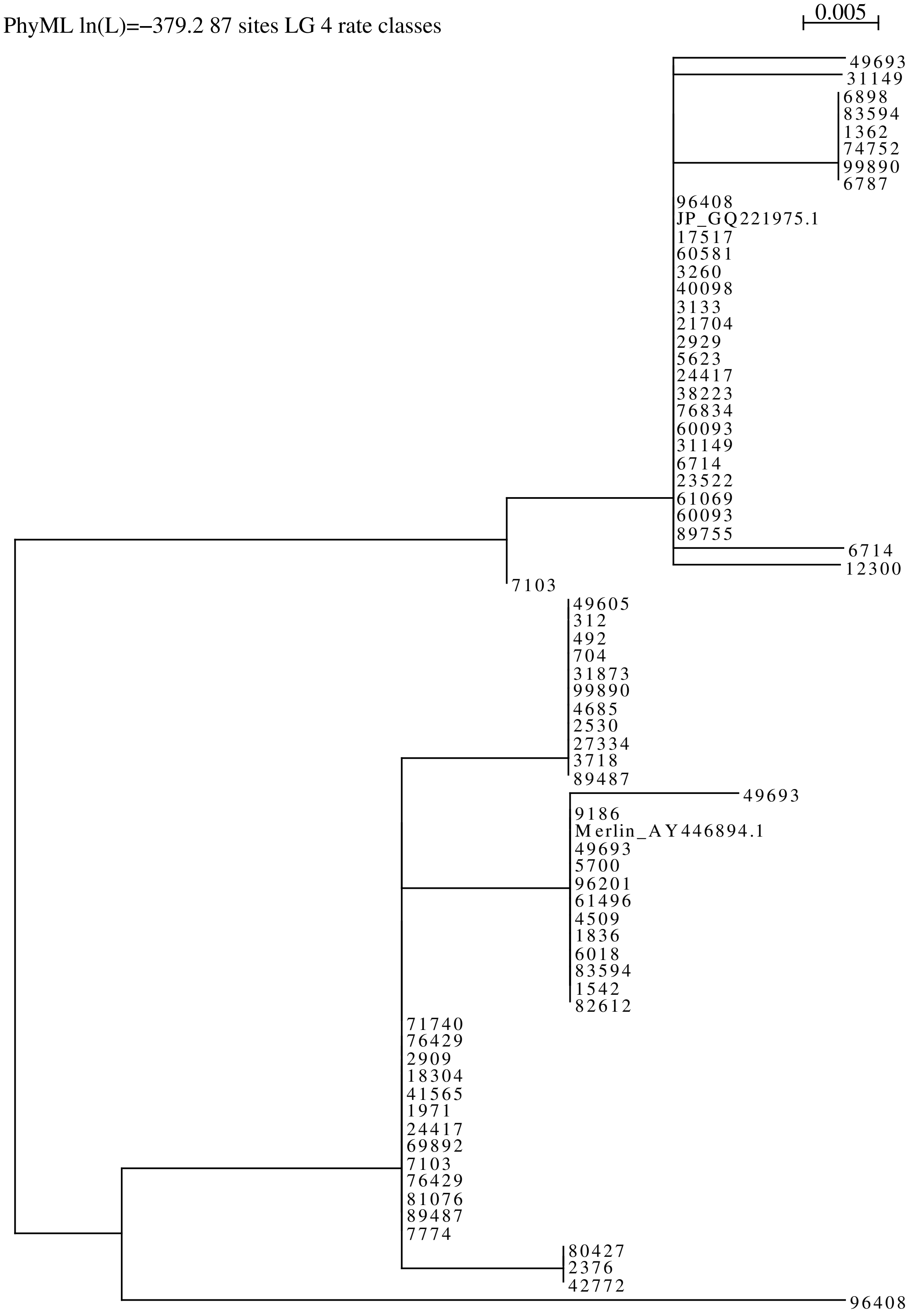

Figure 12: Phylogenetic tree at ORF US27 inferred from the Austrian sample subject to to our article.

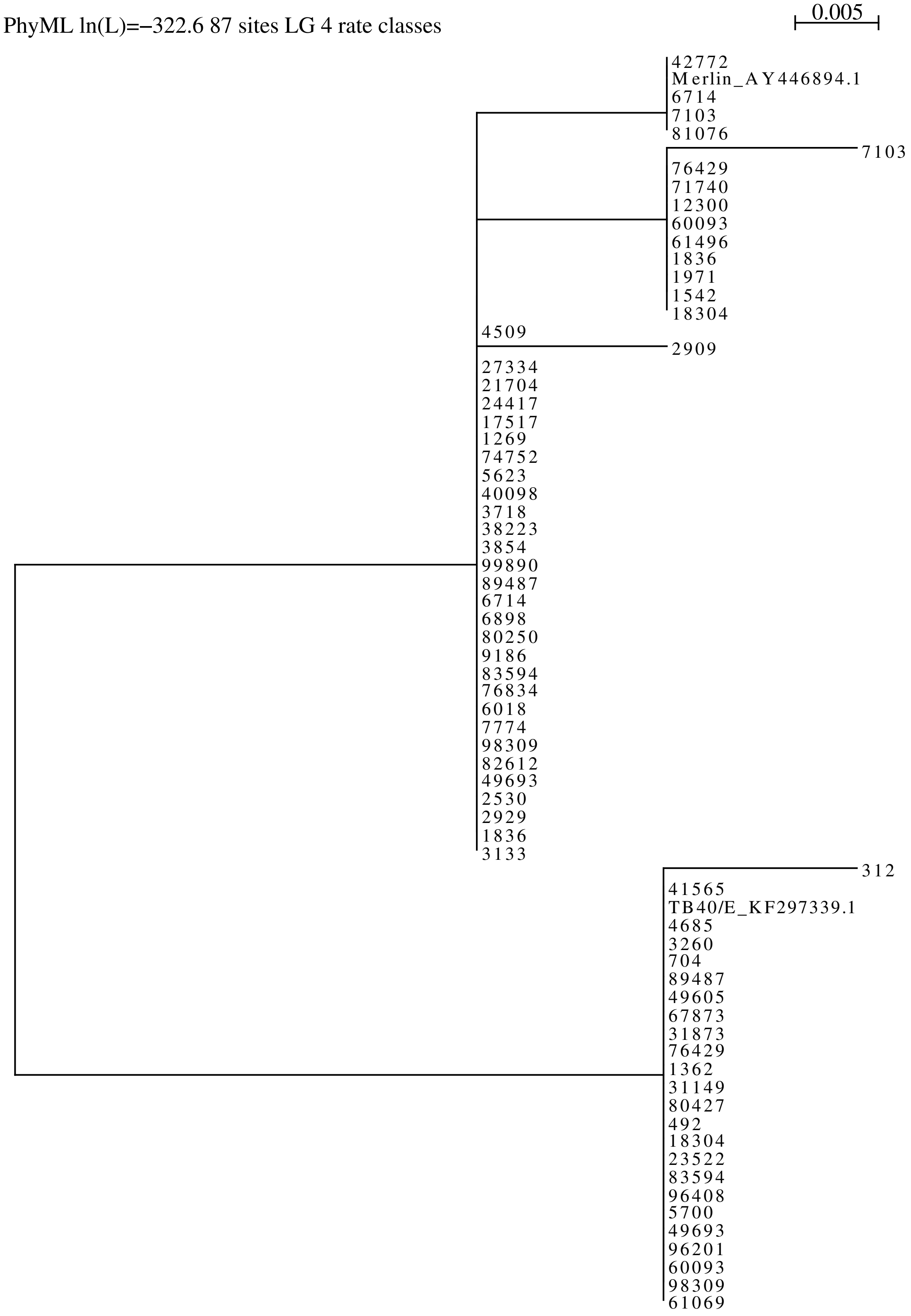
